# Characterization of *Medusavirus* encoded histones reveals nucleosome-like structures and a unique linker histone

**DOI:** 10.1101/2024.04.13.589364

**Authors:** Chelsea Marie Toner, Nicole Marie Hoitsma, Karolin Luger

## Abstract

The organization of DNA into nucleosomes is a ubiquitous and ancestral feature that was once thought to be exclusive to the eukaryotic domain of life. Intriguingly, several representatives of the Nucleocytoplasmic Large DNA Viruses (NCLDV) encode histone-like proteins that in Melbournevirus were shown to form nucleosome-like particles. *Medusavirus medusae* (MM), a distantly related giant virus, encodes all four core histone proteins and, unique amongst most giant viruses, a putative acidic protein with two domains resembling linker histone H1. Here we report the structure of nucleosomes assembled with Medusavirus histones and highlight similarities and differences with eukaryotic and Melbournevirus nucleosomes. Our structure provides insight into how variations in histone tail and loop lengths are accommodated within the context of the nucleosome. We show that Medusavirus histones assemble into tri-nucleosome arrays, and that the putative linker histone H1 does not function in chromatin compaction. These findings expand our understanding of viral histones and suggest that Medusavirus histones represent a snapshot in the evolutionary timeline of nucleosome architecture.

**ONE SENTENCE SUMMARY:** The four *Medusavirus medusae* core histones form nucleosome-like structures that combine features of eukaryotic and other viral nucleosomes.

## MAIN TEXT (INTRODUCTION)

Within the domain of eukaryotes, the compaction of genomic DNA by histones to form nucleosomes is an omnipresent and ancestral feature. The eukaryotic nucleosome core contains four unique histones (H2A, H2B, H3 and H4), each consisting of a structurally conserved histone fold that is common to all four core histones, as well as histone fold extensions (or loops) and highly charged histone tails that are unique to each histone. Two copies each of H2A–H2B and H3–H4 heterodimers assemble into octameric particles that wrap 145-150 bp of DNA to form a canonical nucleosome^1,2^. The histone fold domains of all eight histones are responsible for organizing ∼120 bp of DNA, while the N-terminal α-helix of H3 binds the terminal ∼13 bp of DNA on either side^3^. Histone fold extensions further define the surface of the nucleosome. Numerous protein-protein and protein-DNA interactions within the histone octamer produce a stable disc-shaped particle; however this conformation is more dynamic than originally suggested by the early crystal structures, and this dynamic behavior is essential for the regulatory function of chromatin^1,4^.

The four core histone genes are among the most evolutionarily conserved sequences in the eukaryotic domain of life, suggesting that their incorporation into genomes was an early and essential event during eukaryogenesis^5,6^. There are several hypotheses to explain the enigmatic emergence of the eukaryotic nucleus. Based on the similarities in the information-processing machinery between archaea and eukaryotes, the dominant theory suggests that eukaryotes arose through the metabolic symbiosis of an archaeal host and a proteobacterium, where the gene encoding the single, tail-less histone fold protein of archaea diversified and expanded to give rise to the four core histone genes^5,7–9^. However, the discovery of Nucleocytoplasmic Large DNA Viruses (NCLDV), some of which encode their own viral histone-like proteins, gave rise to the Viral Eukaryogenesis hypothesis. This hypothesis suggests that the early eukaryotic cell was a tripartite conglomerate of an archaeon, an alpha-proteobacterium, and a complex DNA virus (possibly represented by modern NCLDV)^10–12^.

Many NCLDV or ‘giant viruses’ further support this hypothesis by demonstrating various intermediate dependencies on their hosts. They either replicate and assemble in a viral factory that is located in the cytoplasm, or they transiently recruit various host nuclear proteins to the viral factory in the cytoplasm for transcription of early genes^13,14^. Additional evidence supports the hypothesis that the eukaryotic nucleus may be viral in origin, as many NCLDV encode homologues of the critical m7G capping apparatus that is absent in most archaea^15^. In all eukaryotes, the nuclear membrane separates chromatin from the cytoplasm and ribosomes, facilitating the decoupling of transcription from translation with m7G capping. These hallmark eukaryotic genes that are present in the virus represent one critical component of eukaryotic differentiation not widely seen in other domains of life^12^. Therefore, while most giant viral proteins were initially believed to have eukaryotic origin, this alternative hypothesis suggests that they may instead have contributed to modern eukaryotic features. As such, NCLDV may provide insight into the origin of the eukaryotic nucleus^16^.

At least one modern histone-encoding NCLDVs (Melbournevirus, a member of the *Marseilleviridae*) organize their genome in closely packed eukaryotic-like nucleosomes with fused H4-H3 and H2B-H2A doublets^17–19^. *Medusavirus medusae* (MM), named for its ability to turn the *Acanthamoeba castellanii* (*A. castellanii*) host into ‘stone’ through encystment, is one of the few NCLDV known to encode all four core histones (H2A, H2B, H3 and H4) on separate genes. The genome also harbors a putative homologue of the linker histone H1^20,21^. Unlike Melbournevirus, the MM genome enters the host nucleus to initiate DNA replication, while particle assembly and DNA packaging take place in the cytoplasm near the nuclear membrane^20^. The MM putative linker histone H1 is expressed with immediate early genes in the MM life cycle, suggesting a role in reshaping host transcription patterns^20,22^. In contrast, the four core MM histones are expressed at a later timepoint in the infection cycle, suggesting a role in compacting, protecting, and regulating the viral genome, possibly by forming nucleosome-like structures within the established viral factory^20,22^. As such, it is unclear if and where MM core histones are assembled onto viral DNA, and what the role of the putative linker histone is. Compared to eukaryotic and Melbournevirus histones, MM histones differ in the length of histone tails and loops within the histone fold (**Figure 1A, B, S1A-C)**. Additionally, the MM putative linker histone H1 has a dramatically more acidic isoelectric point (pI) than most eukaryotic linker histones (**Figure S1D-F**).

**Figure 1.**
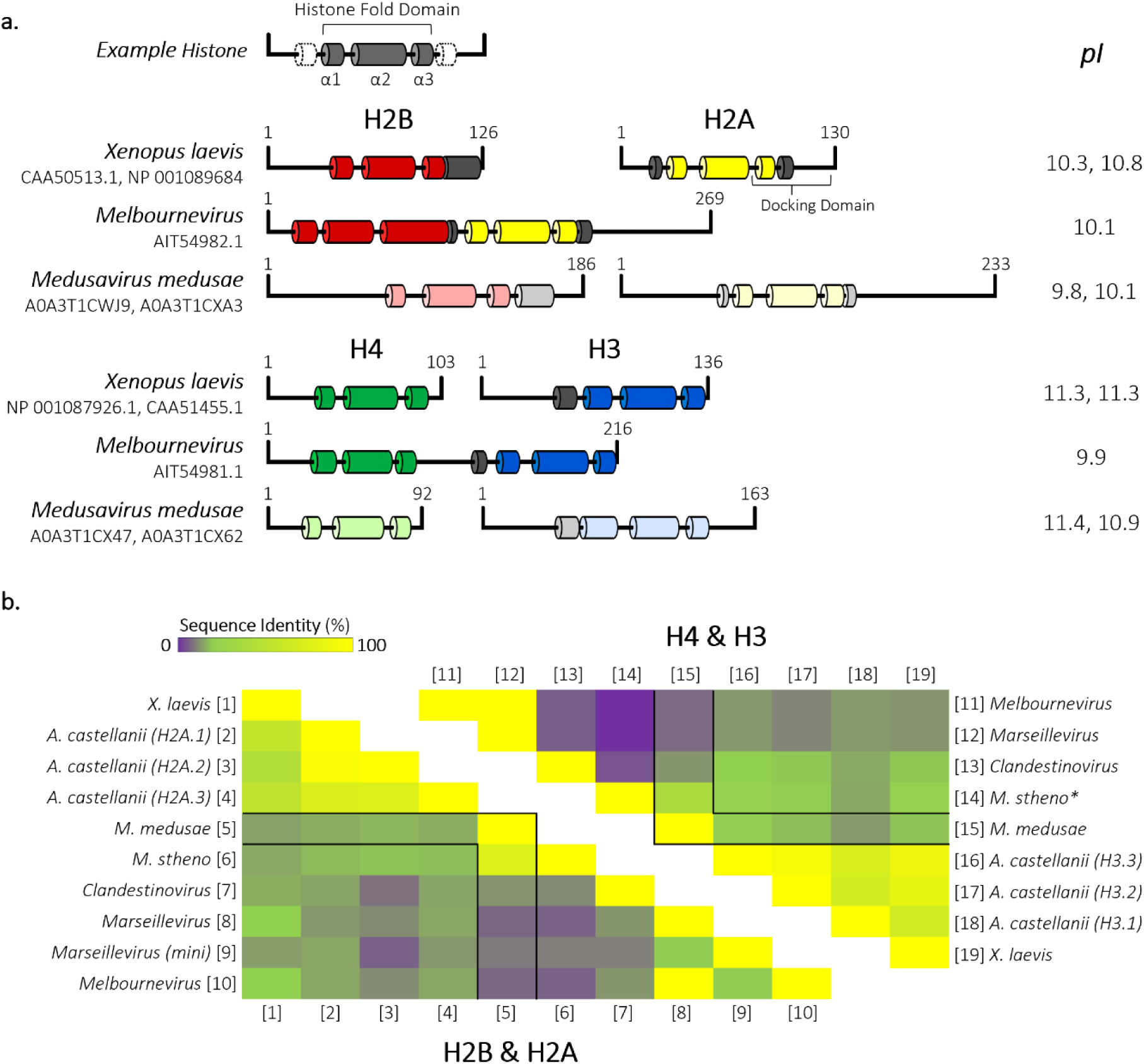
Secondary structure prediction and sequence alignment of *Medusavirus medusae* histones reveals conservation of key eukaryotic residues. (a) Schematic of Eukarya (*Xenopus laevis*) and Nucleocytoviricota histones from Melbournevirus (MV) and Medusavirus (MM). Known *X. laevis* and MV α helices comprising the histone fold domain are represented in dark-colored tubes (H2B, red; H2A, yellow; H4, green; H3, blue; and additional helices, gray). α helices in MM histones were generated using HHpred Quick 2D prediction web server (shown in lighter designated colors). Isoelectric points (pI) of each histone are shown to the right. (b) Heat map comparing percent identity of Eukarya and Nucleocytoviricota H2B-H2A histone sequences (left triangle). Equivalent H2B-H2A histone sequences are represented along the bottom [1-10]. Heat map (right triangle) of percent identity of Eukarya and Nucleocytoviricota H4-H3 sequences. Equivalent H4-H3 histone sequences are represented along the top [11-19]. MM histones are outlined in black within both triangles. A comparison of different dimer pairs (H2B-H2A to H4-H3) sequence identity is not displayed. \**M. stheno* H4-H3 alignment values determined from H3-H4 alignment shown in Figure S1c.

Here, we utilize cryogenic electron microscopy (cryo-EM) to reveal that the predicted core histones from MM form octamers that assemble with DNA into nucleosome-like particles (MM-NLPs). These NLPs are characterized by unique accommodations for elongated loops and tails, and a more pronounced positively charged DNA interacting ridge compared to eukaryotic histone octamers. Additionally, we demonstrate that MM histones can form positioned tri-nucleosomes^23^. AlphaFold and AFM analysis demonstrates that the virally encoded linker histone H1 consists of two winged-helix domains with bimodal charge distribution, but does not promote chromatin compaction, suggesting an alternate virus-specific function. Together, the data presented here underscores the importance of MM-NLPs in the evolutionary timeline of nucleosome architecture and advances our understanding of NCLDV chromatin organization.

## RESULTS

### MM core histones form distinct, stable nucleosome-like particles irrespective of DNA sequence

The MM genome harbors genes for homologs of histones H2A, H2B, H3 and H4 (ORF 318, ORF 61, ORF 255 and ORF 254, respectively)^20^. Secondary structure predictions indicate that MM histones have canonical histone folds (α1–L1–α2–L2–α3), but that the α-helices are connected by longer loops (MM-H2B) or have longer tails (MM-H2B, MM-H2A, MM-H3) than eukaryotic histones (**Figure 1A**). Key eukaryotic nucleosome (eNuc) features are maintained through the H3 αN-helix (which helps organize DNA ends of nucleosomes), but MM-H2A diverges in sequence from eNuc H2A in the docking domain which tethers the H2A-H2B dimers to (H3-H4)_2_ tetramers (**Figure 1A and S1**). MM histones share a 78% sequence identity with histones of its closest relative *Medusavirus stheno,* but less than 30% with eukaryotic histones, and only 23% with the fused histones from *Marseilleviridae* or histones from MM’s closest relative Clandestinovirus. This makes the MM histones unique amongst other known viral histones and more closely related to their amoeba host than other NCLDV (**Figure 1B and S1**). Many signature histone residues are conserved across viral histones including arginine side chains that extend into the DNA minor groove, the paired L1 loops (R-T pairs), and intermolecular histone fold stabilization (R-D clamp). However, MM histones have more predicted DNA-interacting residues in H2A and H2B than in eukaryotic histones (**Figure 1A and S1**).

We expressed, purified, and refolded the four viral histone homologs into an octameric complex (**Figure 2A**). Utilizing the well-established salt-gradient nucleosome reconstitution protocol, increasing amounts of MM octamer were combined with 207 bp DNA (Widom “601” nucleosome positioning sequence) to obtain defined NLPs (**Figure 2B**)^24,25^. We confirmed that MM-NLPs on 207 bp DNA (MM-NLP_207W_) contain a full complement of histones by analyzing sucrose gradient fractions by SDS-PAGE, as well as by analytical ultracentrifugation (see below) (**Figure 2C**, **Table 1, and Figure S2)**.

**Figure 2.**
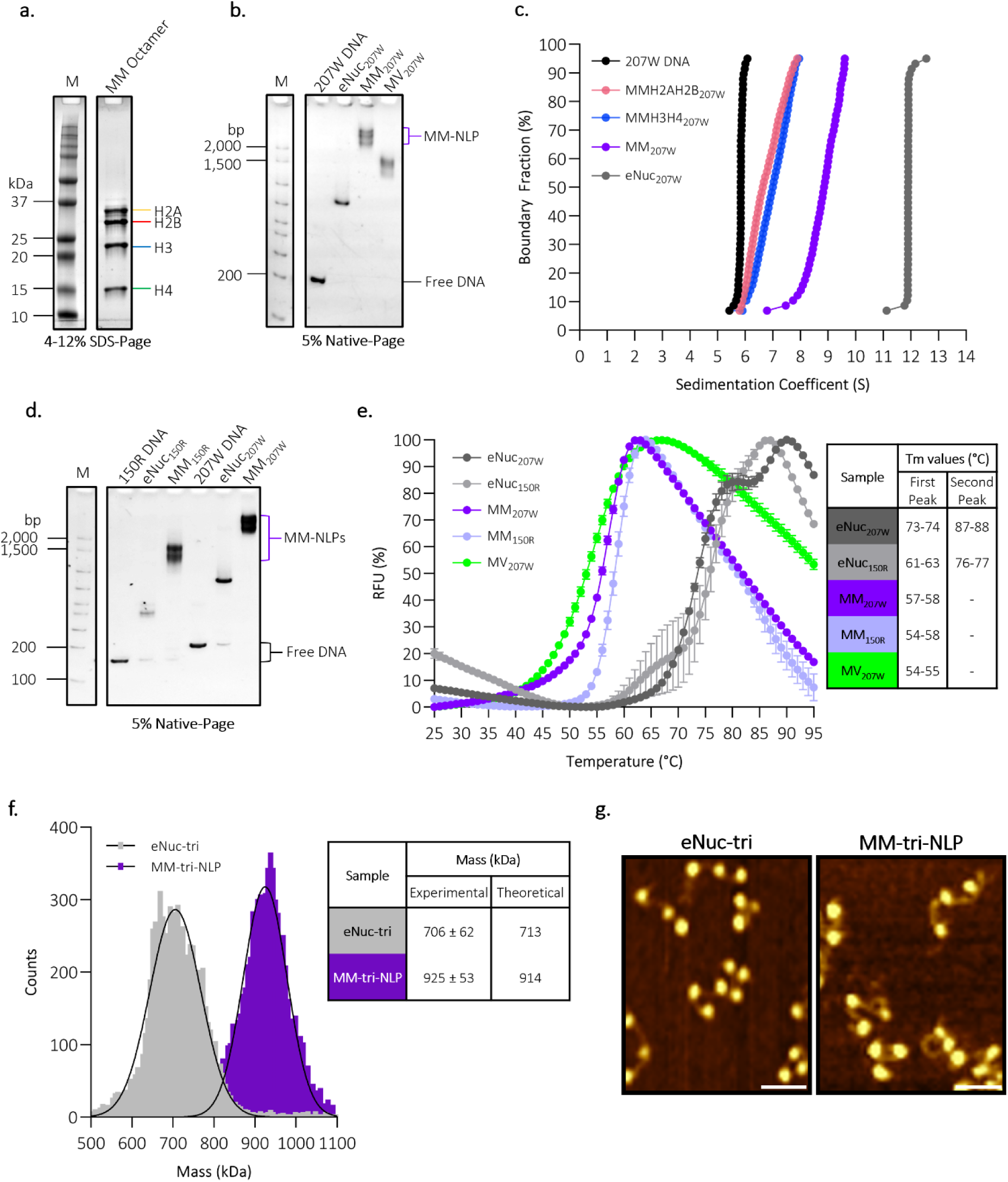
*Medusavirus medusae* histones and DNA assemble into stable mono nucleosome-like particles (NLP) and tri-NLP *in vitro*. (a) SDS-PAGE of octameric refolded *Medusavirus medusae* (MM) core histones H2A, H2B, H3, and H4. (b) MM-NLP, eNuc, and Melbournevirus-NLP (MV-NLP) reconstituted with Widom ‘601’ 207 bp DNA and analyzed by 5% Native-PAGE stained with SYBRGold (DNA visualization). (c) SV-AUC of reconstituted NLP and histone-DNA complexes. Van Holde-Weischet plot of eNuc on 207 bp DNA, histone-DNA complexes with H2A-H2B and (H3-H4)_2_, and MM-NLP207. Quantitative evaluation is given in Table 1. (d) 5% Native-PAGE of reconstituted MM-NLP and eNuc on 150 bp ‘random sequence DNA’ (50% G/C DNA (150R)), and on Widom ‘601’ 207 bp DNA. (e) Thermal stability of MM-NLPs and eNucs shown in panel (d) including MV_207W_ from panel)b) (n=2). Averages of the replicates are shown and normalized with standard deviations denoted. Tm values of each MM-NLP and eNuc are shown in the inset. (f) Mass photometry analysis of eNuc-tri (gray) and MM-tri-NLP (purple). The solid lines represent the Gaussian function fit to the main species observed on particle counts versus molecular mass distribution histograms, with the estimated molecular weight (in kDa) corresponding to the respective mass at the center of each peak. Theoretical and measured molecular masses (± SD) are shown in the inset. (g) Representative AFM topography images of eNuc-tri and MM-tri-NLP. Scale bar = 50 nm. Full images are shown in Figure S3C and D.

**Table 1.**
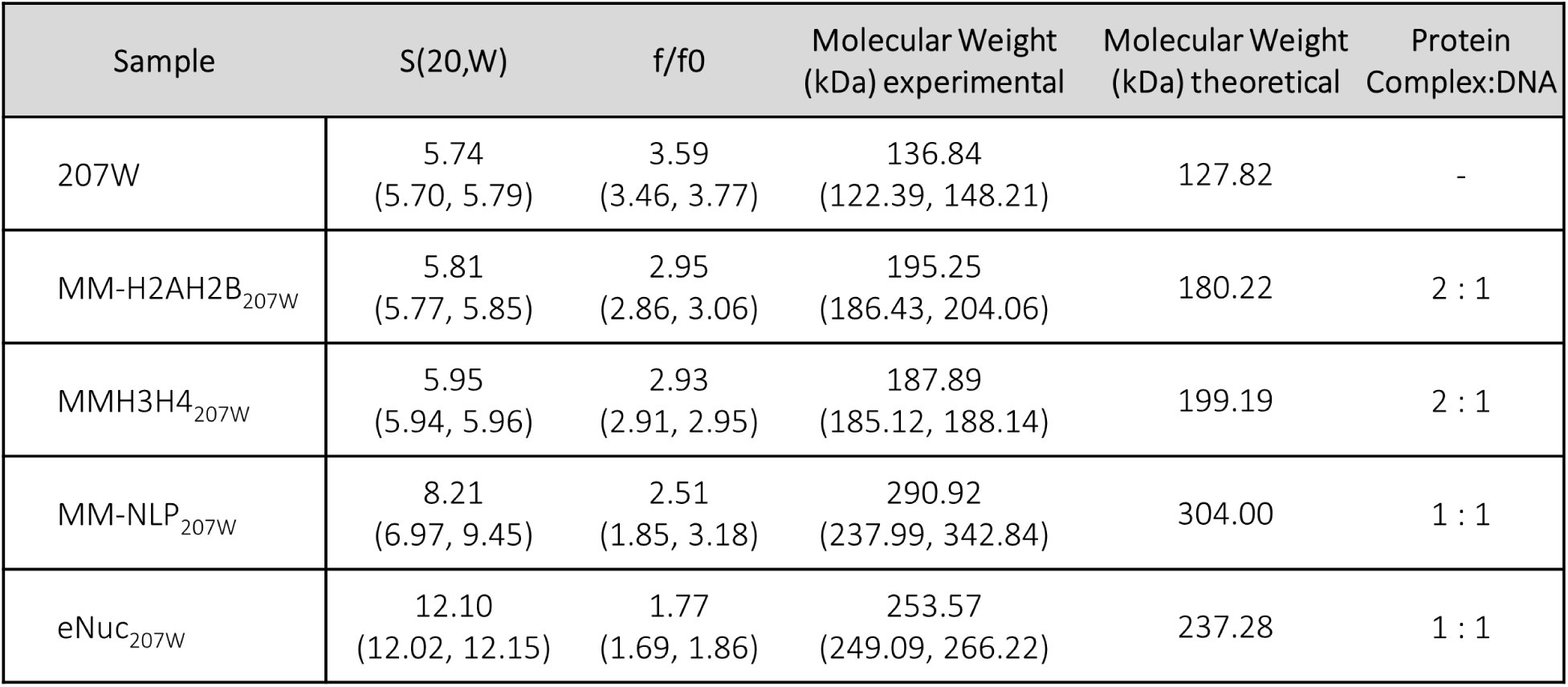
S values (S(20,W)), frictional ratios (f/f0), and calculated molecular weights (with confidence intervals) of histone-DNA complexes from SV-AUC. Theoretical protein complex: DNA ratios given are calculated for H2A-H2B dimer, (H3-H4)2 tetramer, and histone octamer. All values were calculated using UltraScan ^52,53^.

MM-NLPs migrate much higher on native gels compared to eukaryotic nucleosomes (eNuc), and even higher than Melbournevirus NLPs. Sedimentation velocity analytical ultracentrifugation (SV-AUC) allows the determination of macromolecule size and overall shape in solution from diffusion-corrected sedimentation values, which are proportional to particle mass and inversely proportional to viscous drag^26^. eNuc_207W_ sediments at 12 S, comparable to previous eNuc_147W_ sedimentation at 11 S^17^. MM-NLP_207w_ sediments at ∼ 8 S. This decrease is likely caused by an increased viscous drag compared to eNuc, and is similar to what was observed for Melbournevirus NLP_207w_ (**Table 1**). Reconstituting either MM H2A-H2B dimer or MM (H3-H4)_2_ tetramer onto the same DNA gives rise to particles with the expected molecular weight and an even higher level of viscous drag (f/f_0_). Importantly, the experimentally determined molecular mass of MM-NLP_207w_ (290.92 ± 51.92 kDa) is in close agreement with the expected theoretical mass of an octameric protein complex with 207 bp DNA (304.00 kDa) (**Figure 2C and Table 1**).

To determine if MM-NLP formation is affected by either DNA sequence or length, we reconstituted MM octamer with a 150 bp ‘random’ DNA sequence (MM-NLP_150R_). This DNA was designed to have a 50% GC content over each 10 bp segment and does not harbor strong nucleosome positioning signals. Both MM-NLP_150R_ and MM-NLP_207W_ migrate much higher than eNuc on native-PAGE, supporting the finding that MM histones form less compact particles irrespective of DNA sequence (**Figure 2D**).

The stability of the nucleosomes and their corresponding histone octamers was tested in a thermal melting assay (25 °C – 95 °C) by monitoring the fluorescence of SYBRO Orange release from denatured complex. The MM octamer fluoresces at 25 °C, indicating instability, as opposed to the eukaryotic octamer that does not begin to dissociate until ∼40 °C (**Figure S3A**). eNucs on both DNA fragments (207W and 150R) demonstrate the characteristic peaks of histone dimer and tetramer release from the nucleosome, as previously reported^27^. In contrast, all viral NLPs (MM-NLP_150R_, MM-NLP_207W_, and Melbournevirus NLP_207W_) melt in a single peak at much lower temperatures, suggesting the dissociation of the octamer from DNA in a single step, and irrespective of sequence and length. This demonstrates lower thermal stability of viral octamers and nucleosomes compared to their eukaryotic counterparts, as previously shown for Melbournevirus NLP (**Figure 2E**)^17^.

To test if MM-NLP can assemble into higher-order chromatin, we reconstituted MM tri-nucleosome-like particles (MM-tri-NLP) onto a 3x copy of Widom 207 DNA^28^. As observed for the single MM-NLP, MM-tri-NLPs migrate higher on a native gel relative to *X. laevis* tri-nucleosomes (eNuc-tri) (**Figure S3B**). The mass of the MM-tri-NLP was estimated in solution by mass photometry to be 925 kDa ± 53 kDa, highlighting the considerable size difference of the viral histones compared to eNuc-tri at 706 ± 62 kDa (**Figure 2F**). MM-tri-NLPs exhibit the same characteristic “beads on a string” structure as eukaryotic tri-nucleosomes, forming three distinct particles as visualized by Atomic Force Microscopy (AFM) (**Figure 2G).** Consistent with the decreased stability of MM-mononucleosomes, the MM-tri-NLP appear more heterogenous than eNuc-tri in AFM analysis, likely due to the presence of nucleosomal sub-species during salt gradient dialysis or disassociation during dilution **(Figure S3C and S3D**).

### MM-NLP resemble eNuc with unique accommodations for longer histone tails and loops

Single-particle cryogenic electron microscopy (cryo-EM) was utilized to determine the structure of MM-NLP_207W_. Samples were crosslinked through gradient fixation (GraFix) with glutaraldehyde^17^, and compared to a native sample that had not been subjected to crosslinking. The migration in the sucrose gradient was unchanged upon crosslinking (**Figure S2**). Data for both particles was collected on a Titan Krios G3i, and following 3D reconstruction and refinement, we obtained an electron density map of crosslinked MM-NLP_207W_ at 4.3 Å resolution (**Figure 3 and S4A**), and at 5.1 Å resolution for native MM-NLP_207W_ (**Figure S4B**). We observed electron density for all core histone helices and ∼ 135 bp of DNA, allowing assignment of the MM-NLP_207W_ core. Density for the MM-H2A tail region of the docking domain was not visible. Our initial homology model of MM histones in a nucleosome-like arrangement was refined by docking into the MM-NLP_207W_ density, with the finalized model demonstrating strong agreement (cross correlation=0.705) (**Figure 3A**).

**Figure 3.**
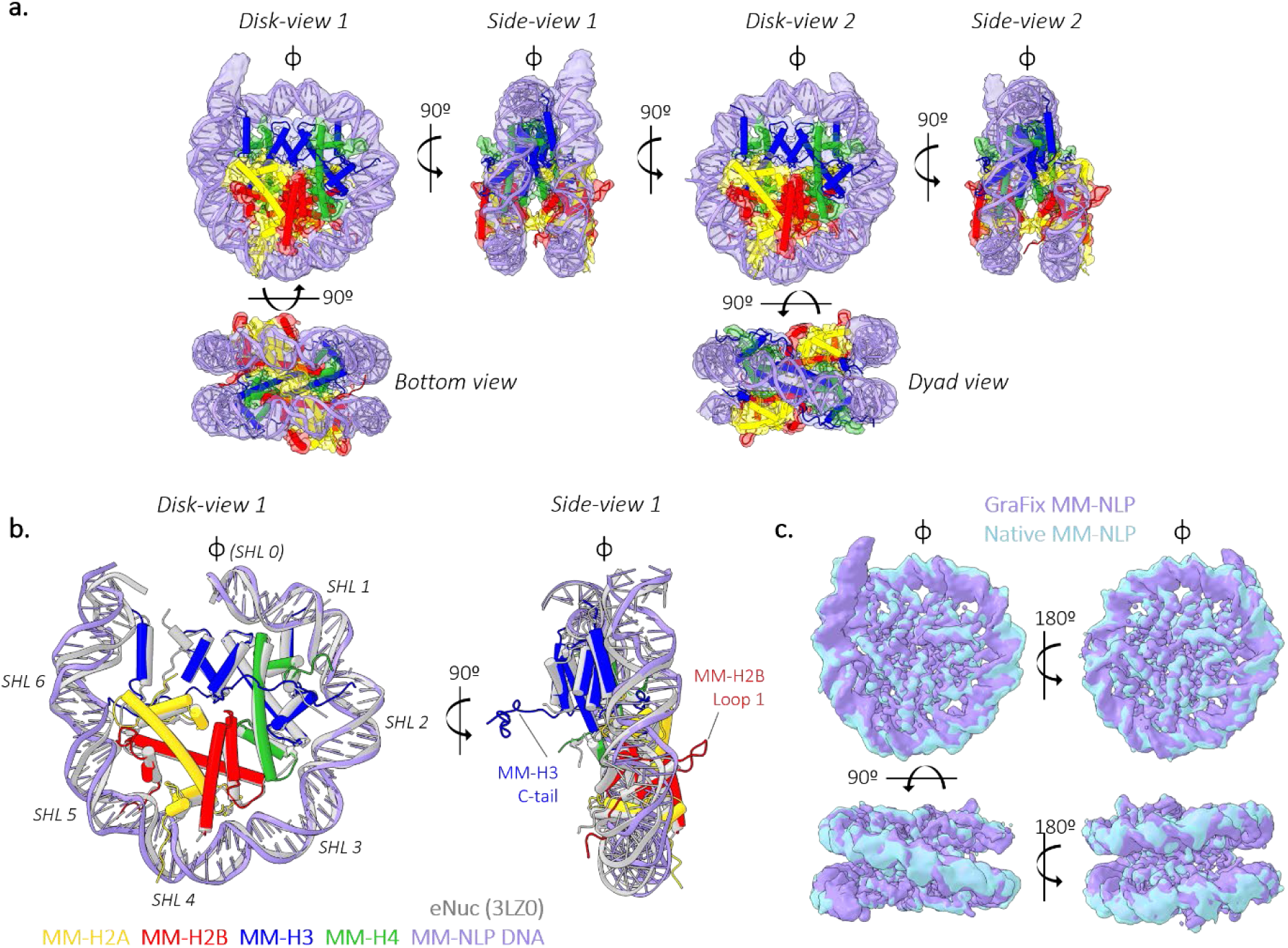
*Medusavirus medusae* NLP (MM-NLP_207W_) closely resemble eukaryotic nucleosomes. (a) Overview of MM-NLP_207W_ structure. Individual histones H2A, H2B, H3 and H4 and their surrounding density are represented by yellow, red, blue, and green respectively. (b) Overlay of MM-NLP207 (H2A-yellow, H2B-red, H3-blue, H4-green, DNA-purple) with eNuc (gray) with only 71 bp of DNA, one H2A-H2B dimer, H3-H3’ four helix bundle, H3’ N-terminal helix, and a single H4 displayed for clarity. Superhelix locations (SHLs) are numbered from 0 to 6 starting at the nucleosome dyad (ɸ). (c) Comparison of native (light blue) and GraFix (purple) MM-NLP207 electron densities.

Just like its eukaryotic counterpart, the MM-NLP_207W_ contains two copies each of H2A, H2B, H3 and H4 as an octameric core formed by histone fold regions, wrapped by ∼ 130 bp of DNA (**Figure 3A**). However, the density of only one of the two H3 αN-helices and associated DNA is observed with confidence, underscoring the dynamic character of the ∼13 penultimate base pairs of DNA which has been noted in several eukaryotic and Melbournevirus nucleosome structures^29,30^. The overall geometry of the DNA superhelix and the layout of histone fold helices are near-identical between MM-NLPs and the eukaryotic nucleosome, with minor differences in the wrapping of DNA ends (**Figure 3B**). The structure of native (i.e. not crosslinked) Melbournevirus NLP_207W_ at 5.1 Å has a high correlation (0.949) with crosslinked Melbournevirus NLP_207W,_ confirming that there were no induced and potentially artificial conformations due to crosslinking (**Figure 3C and 4A)**.

**Figure 4.**
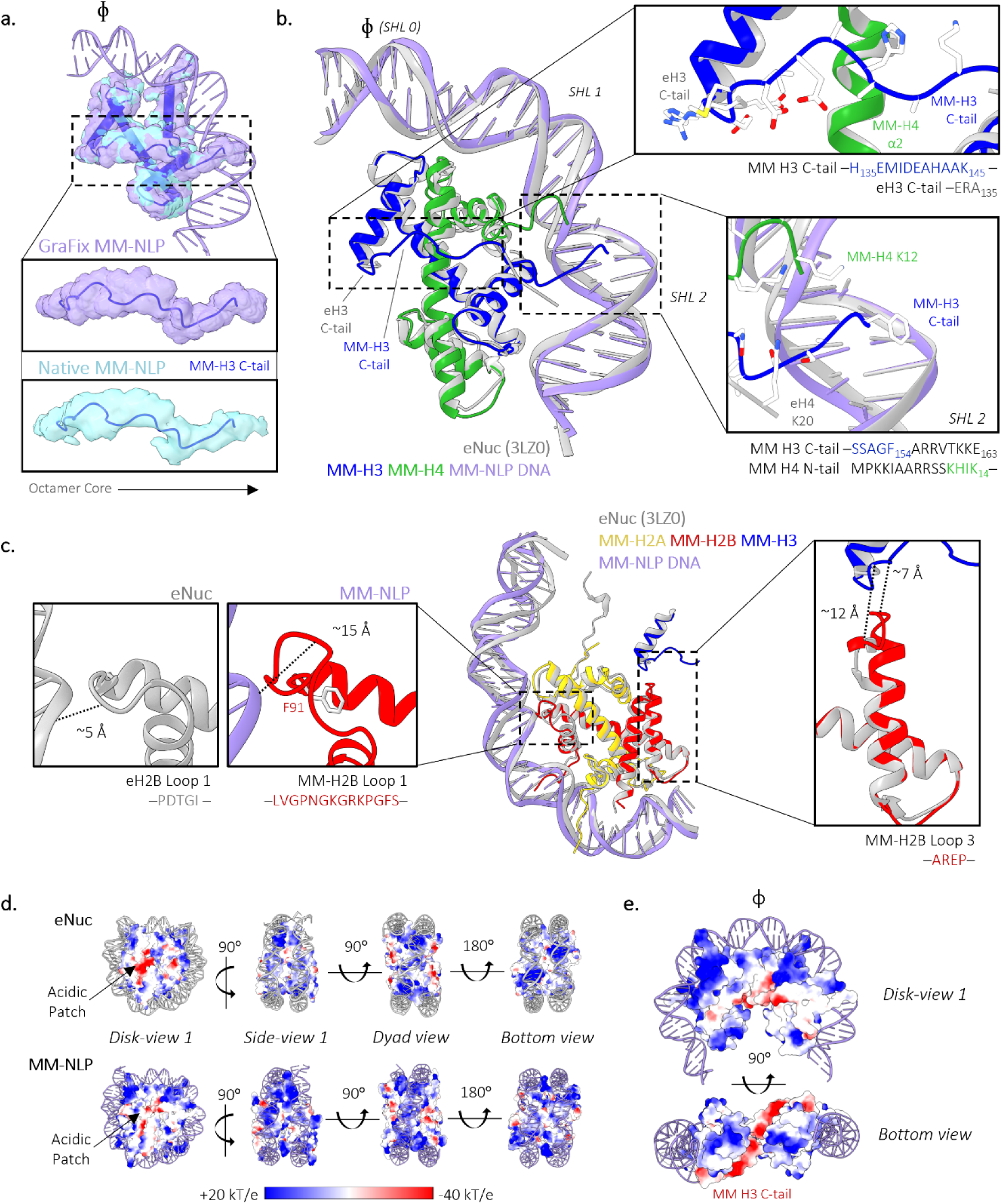
Unique roles and accommodations for longer tails and loops in MM-NLP. (a) Superposition of GraFix MM-H3 density and native MM-H3 density. The MM-H3 tail from H135 to F154 is displayed in the inset. (b) Superposition of MM-(H3-H4) with eukaryotic (H3-H4) and 30 bp of associated DNA. Close-ups are provided of MM H3 C-tail orientation in MM-NLP in relation to eukaryotic H3 and the MM-H3 C-tail extension to SHL 1.5. Corresponding residues of each tail are denoted below each close-up, with tail end residues that are not resolved shown in black. (c) Superposition of MM-H2A-H2B dimer aligned with eukaryotic H2A-H2B dimer (light gray) and 40 bp of associated DNA. Close-ups are provided of both eH2B and MM-H2B loop 1 along with the MM-H2B α3 and loop 3 extension towards the MM-H3 C-tail. Distances were measured from the backbone atoms of the closest residues from each loop to DNA or H3 C-tails. (d) Charged surface representation of histones from the MM-NLP_207_ and eNuc (PDB ID: 3LZ0). (e) Charged surface representation of MM (H3-H4)_2_ tetramer with 60 bp of DNA.

MM histones have distinctly longer tails (H2A, H2B, H3) and loops (H2B) than eukaryotic histones. Most unusually, MM-H3 has a 29 amino acid long C-terminal tail that is not observed in any other H3 histones **(Figure 3B, S1B)**. Both of our structures (crosslinked and native) reveal that this H3 tail extends from the end of α3 and lays across its partnered H4 α2-helix to reach the DNA minor groove at SHL ±1.5 (**Figure 4A and 4B**). In this position, it is in close contact with the H2A-H2B dimer, H4 α2, the H4 N-terminal tail, and H3 α1 through a combination of hydrophobic packing and hydrogen bonding interactions. The density for this tail is very well-defined and is observed in both the crosslinked and native structure, excluding crosslinking artifacts. The presence of the extended H3 C-terminal tail redirects the MM-H4 N-terminal tail to contact the DNA at SHL ± 1.5 (**Figure 4B**). The penultimate 10 amino acids of the H3 C-terminal tail (containing three arginines and two lysines) are poised to interact with SHL ± 5.5 but are too disordered to be observed in the maps.

A second unique feature of MM histones is the extended H2B L1 loop connecting α1 and α2. This loop, which is 11 amino acids longer compared to eukaryotic histones and resembles a β hairpin, has very well-defined density that protrudes from the nucleosome disk by about ∼ 15 Å (**Figure 3B**)^31^. Its base packs against H2B α2-L2 by forming a hydrophobic core centered around H2B F91 (**Figure 4C**). Together with α1, this extended loop forms a defined module that contributes to stabilizing the interaction with H2A, and uniquely defines the surface of the MM-NLP. Additionally, the α3 helix of MM-H2B is longer which allows the extended connector (4 amino acids) between H2B α3 and αC to contact the H2A docking domain and the extended H3 C-terminal tail, contributing to their unique interactions within the histone octamer (**Figure 4C**).

MM-NLPs exhibit a more pronounced positively charged electrostatic DNA-interacting ridge compared to eNuc. Full views of the ridge show a higher density of positive charge in MM-NLP, consistent with the predicted increase in DNA-interacting residues (**Figure 4D**, **Figure 1A, Figure S1**). The acidic patch, a localized region on each disk “face” of the nucleosome, is a well-established binding site for many chromatin-interacting proteins in eukaryotes^32^. The negative charge in this region of the MM-NLP is less pronounced (**Figure 4D**), possibly due to our inability to build parts of the H2A docking domain. Notably, the H3 C-terminal tail contributes to a pronounced “S” shaped acidic surface along the bottom side of the (H3-H4)_2_ tetramer that is unique to MM histones (**Figure 4E**).

### Medusavirus NLPs are structurally more similar to eukaryotic nucleosomes than to other viral NLPs

To date, the only other viral NLP for which structural information is available is the Melbournevirus nucleosome, where histones are fused into doublets (H2B-H2A and H4-H3)^17,19^. Overall, the two structures are rather similar to each other in the positioning of the histone fold elements and the DNA. An intriguing commonality between the viral NLPs is the presence of a histone tail that lays across the same region of the H4 α2-helix (**Figure 5A**). In MM-NLP, as shown above, the extended H3 C-terminal tail (unique to MM-H3) reaches over H4 α2, while in Melbournevirus the redirected N-terminal tail of H4 assumes the same function in but coming from the opposite direction. The tails contact similar positions on top of the H4 α2-helix (centered around H4 V50) (**Figure 5B**). The interface consists of charge-charge interactions and hydrophobic packing between the H4 α2-helix and the respective tail, signifying an intriguing functional convergence between the two distantly related viral nucleosome particles (**Figure 5B**).

**Figure 5.**
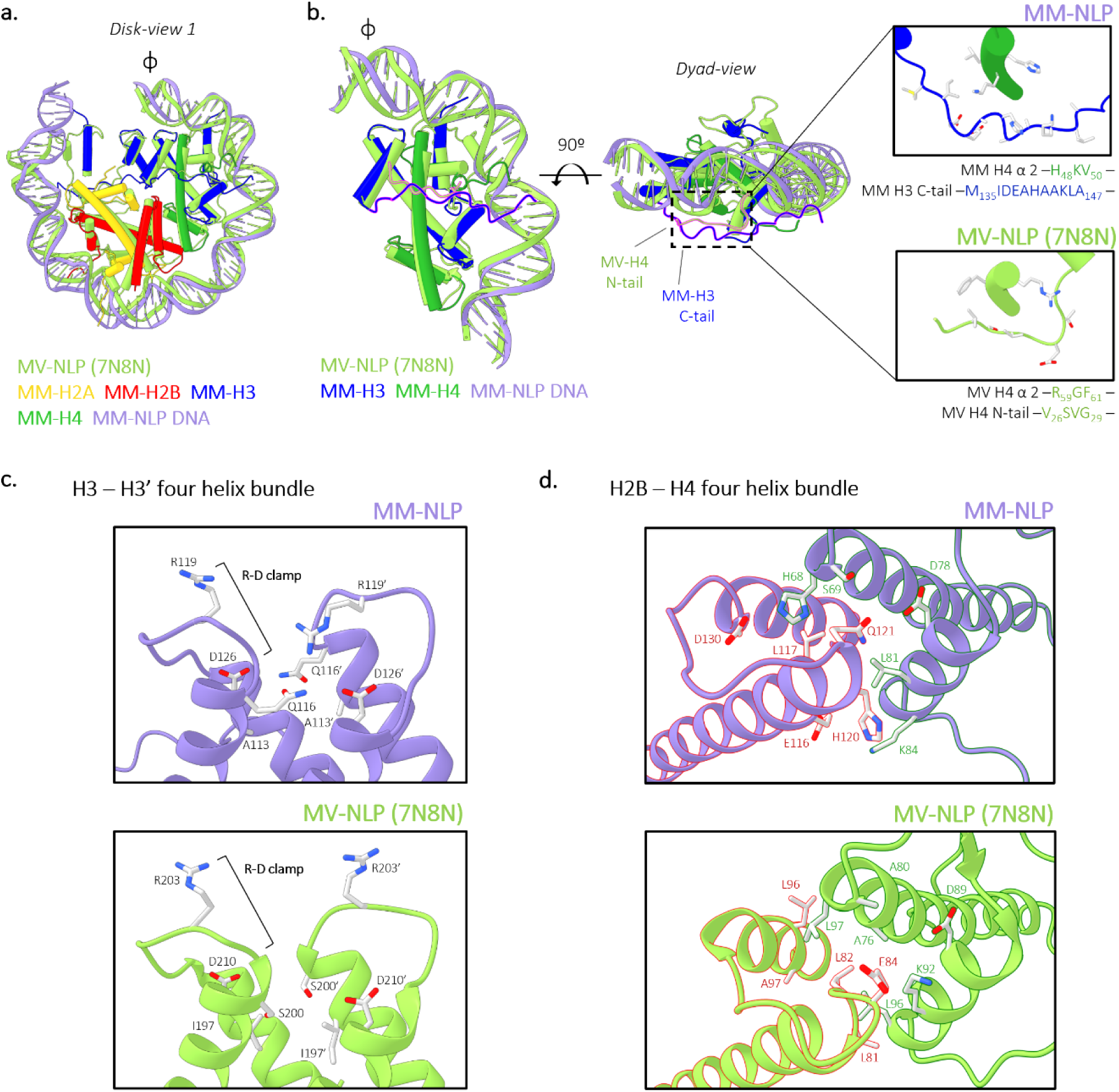
Structural comparison of viral nucleosomes exposes distinct accommodations and variations in intermolecular interactions of viral NLP. (a) Superposition of MM-NLP and Melbournevirus-NLP (MV-NLP; light green; PDB ID: 7N8N). (b) Superposition of H3-H4 histones of MM-MVP and H4-H3 histone doublet of MV-NLP (light green). MM-H3 C-tail and MV-H4 N-tail are outlined in pink. Inset highlights the MM-H3 C-tail and MM-H4 N-tail pathway across the H4 α2 of each viral NLP. Residues within the inset are denoted below each close-up. (c) Residues contributing to the H3 – H3’ four helix bundle interactions of MM-NLP and MV-NLP. The stabilizing R-D clamp seen in eukaryotes is highlighted in both insets. (d) Residues contributing to the H2B – H4 four helix bundle interactions of MM-NLP and MV-NLP.

The charge distribution of the histone octamer surface differs between the two viral NLPs, with MM-NLP displaying a significantly more positively charged DNA interacting ridge compared to Melbournevirus NLP **(Figure S5**). Superposition of MM-NLP and Melbournevirus NLP with eNuc allows us to highlight conserved histone-histone and histone-DNA interactions. Key features such as the H3 ‘R-D clamp’ and the ‘sprocket arginine’ from H2A are present in MM-histones, and as such are conserved between viral and eukaryotic NLPs (**Figure 5C**). However, MM-NLP maintains the arginine minor groove interactions from H3 and H4 while Melbournevirus NLP does not have the equivalent residues.

The histone core is held together by four-helix bundle interactions between H3 and H3’, and between H2B and H4. MM-NLP differs in both of these interfaces from Melbournevirus NLP (**Figure 5C and 5D**) and eNuc. In MM-NLP, both interfaces are characterized by glutamine residues, and are less hydrophobic in nature than either Melbournevirus NLP or eNuc. Both viral nucleosomes lack the histidine-cysteine configuration in the H3-H3’ four-helix bundle that is typical to nearly all eukaryotic nucleosomes, and that is hypothesized to convey copper reductase activity^33^.

### MM putative linker histone H1 does not compact tri-nucleosomes

Unique amongst histone encoding NCLDVs, MM encodes a putative linker histone H1 (ORF 106) with an uncharacteristically acidic pI of 5, similar to the pI of H1.1 of its host *A. castellanii* (**Figure 6A and S1E**)^20^. The putative MM linker histone H1 (MM-H1) has a low sequence conservation (12.86 %) compared to eukaryotic *X. laevis* H1.0 and *Gallus gallus* H5. Secondary structure predictions of MM-H1 suggest the presence of the canonical N-terminal winged-helix DNA-binding domain (**Figure S1D**). Unexpectedly, we predict an additional winged-helix domain in both MM-H1 and in all *A. castellanii* H1 variants following a variableloop, which is supported by AlphaFold structures (**Figure 6A and S6**). Both winged-helix domains predicted for MM-H1 have a more negative charge compared to *X. laevis* H1.0, with the second predicted domain displaying a predominantly negative charge (**Figure S6**). Superimposing MM-H1 with *X. laevis* H1.0 reveals that the four basic amino acids (K40, R42, K52 and R94) required to compact chromatin are not conserved in MM-H1^34^. Instead, the predicted MM-H1 has one equivalent basic residue (R69) and acidic residues (D18, P20 and D30), which suggests MM-H1 may lack the ability to function in compaction (**Figure 6B**).

**Figure 6.**
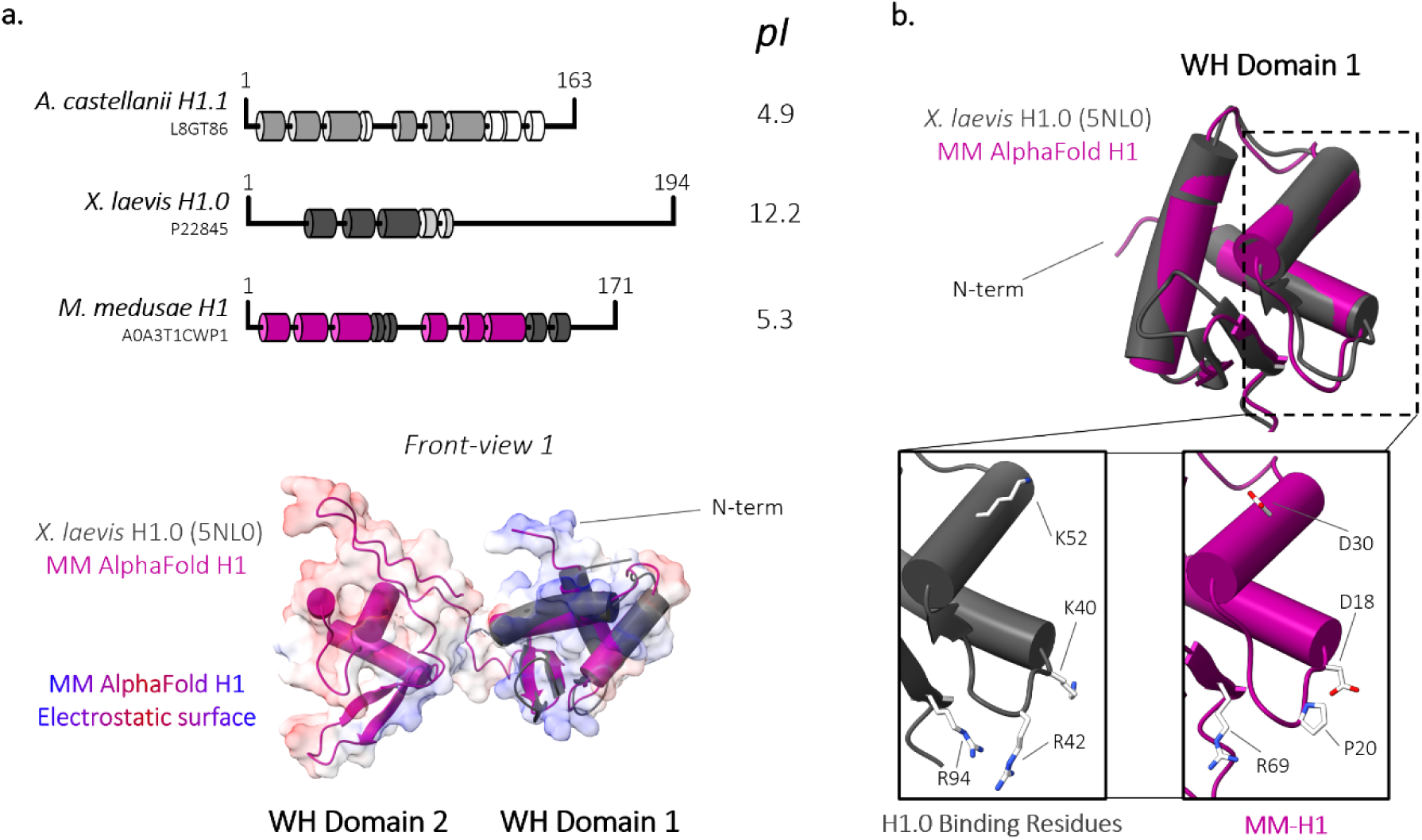
*Medusavirus* linker histone H1 contains a second winged-helix domain and lacks canonical compaction residues. (a) *Acanthamoeba castellanii* (*A. castellanii*) H1.1, *Xenopus laevis* (*X. laevis*) H1.0, and *Medusavirus medusae* (MM) linker histones were aligned using HHpred’s multiple sequence alignment tool (ClustalΩ). α helices and β sheets were predicted using HHpred’s Quick 2D prediction web server (shown in dark pink). Known *X. laevis* H1.0 and *A. castellanii* α helices and β sheets representative of the winged helix (WH) domain is represented in dark-grey tubes and light-grey tubes, respectively. Corresponding isoelectric points (pI) of each linker histone are provided to the right. Superposition of AlphaFold MM-H1 (dark pink) and *X. laevis* H1.0 (dark grey; 5NL0) within the charged surface representation of MM-H1. (b) Overlay of MM-H1 (shown in dark pink) and *X. laevis* H1 (dark grey; 5NL0). Inset: *X. laevis* H1 residues involved in chromatin compaction and corresponding residues in MM-H1.

To characterize the putative MM linker histone H1 (MM-H1), we expressed, purified, and refolded the protein for biochemical analysis (**Figure S7A**). Circular dichroism of refolded MM-H1 confirms a significant increase in order when compared to *Mus musculus* H1.0 (eH1.0), from ∼35% to ∼75% (**Figure S7B and S7C)**, consistent with the predicted presence of a second winged-helix domain in MM-H1 (**Figure 6**)^35,36^.

AFM has previously been used to demonstrate eNuc-tri compaction by the linker histone eH1.0, which was characterized by an increase in particle height, as well as by an increase of visually compacted particles^37^. We recapitulated eNuc-tri compaction with the addition of eH1.0 by observing an increase in the height distribution of tri-nucleosome particles (**Figure 7A, 7B and 7E**). This is expected based on structures of H1 bound nucleosomal arrays, which show a zig-zag arrangement with non-consecutive nucleosomes 1 and 3 forming a stack, and H1 bound to linker DNA near the dyad of each nucleosome^38^. This stacking likely explains the “bi-lobed” tri-nucleosomes we observe via AFM when tri-nucleosomes are incubated with eH1.0. The stacked terminal nucleosomes (nucleosome 1 and 3) have increased height compared to the central nucleosome (nucleosome 2) (**Figure 7A, 7B, and 7E**). The addition of MM-H1 to eNuc-tri did not yield these distinct “bi-lobed” tri-nucleosomes or an increase in particle height (**Figure 7C, 7D, and 7E**). Compaction was also directly assessed by measuring the total distance between the three nucleosomes in the tri-nucleosomes. Upon the addition of eH1.0 to eNuc-tri there was a significant decrease in total distance between nucleosomes that was not seen upon the addition of MM-H1 (**Figure 7F**). However, this analysis precludes the ability to measure the most compact tri-nucleosomes where nucleosomes are not individually visible (i.e. bi-lobed), thus underestimating the level of compaction observed with eH1.0. Using these two AFM analyses, no tri-nucleosome compaction was observed upon the addition of MM-H1. Together, this data suggests that MM-H1 does not function in compaction under these conditions, as predicted by the lack of key compaction residues and acidic electrostatic surface (**Figure 6B and S6**).

**Figure 7.**
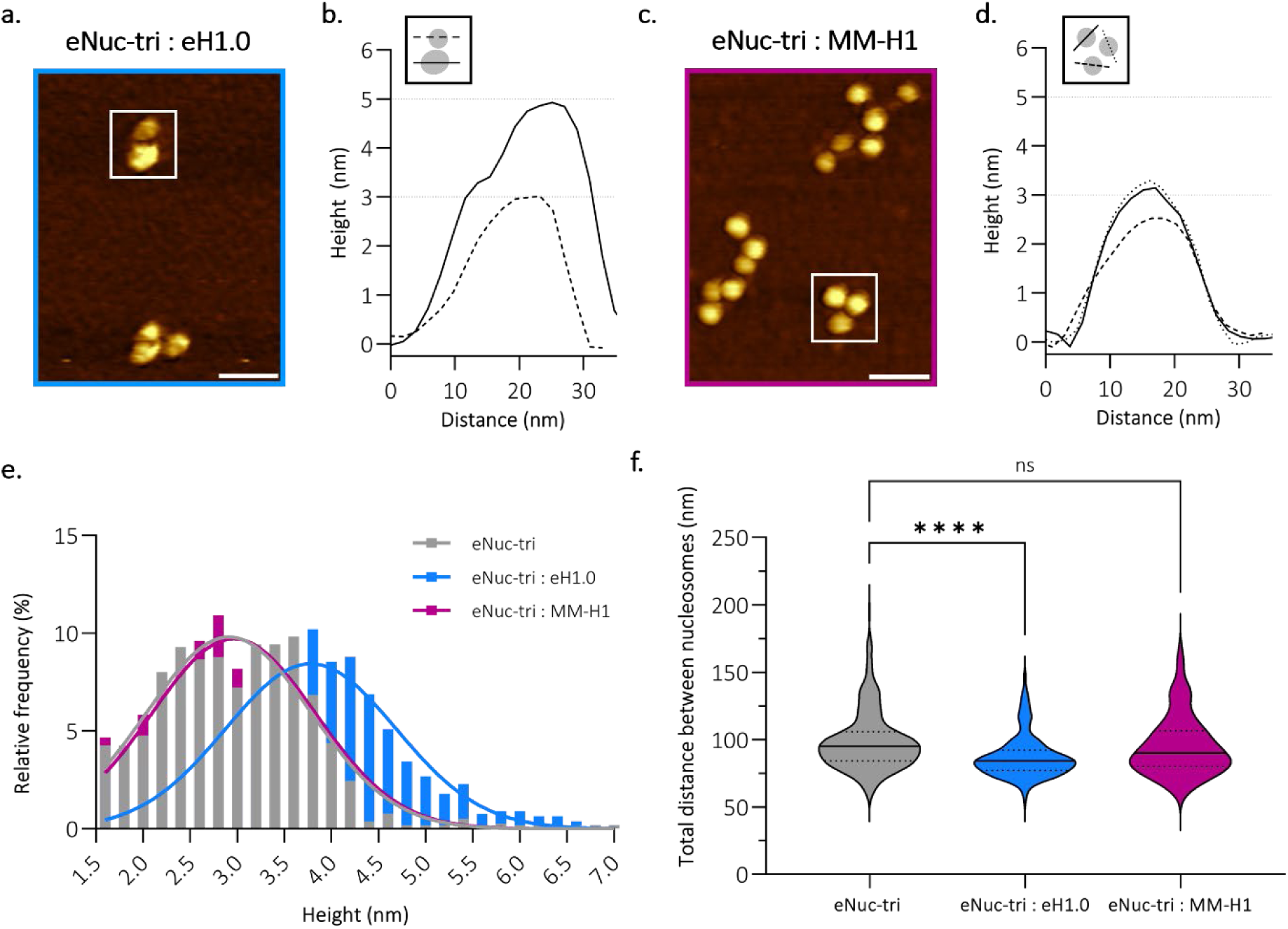
*Medusavirus medusae* linker histone H1 does not compact tri-nucleosomes. (a) Representative AFM topography image of eNuc-tri:eH1.0 sample, imaged in air, with (b) height profile along the line through indicated particles shown in graphical inset. (c) Representative AFM topography image of eNuc-tri:MM-H1 sample, imaged in air, with (d) height profile along the line through indicated particles shown in graphical inset. Dashed lines at 3 and 5 nm shown for reference. Scale bar = 50 nm. (e) Histogram and Gaussian fitting of particle height for eNuc-tri alone (2.9 ± 0.9 nm, *N* = 774, grey) and after incubation with either eH1.0 (3.9 ± 0.9 nm, *N* = 784, blue) or MM-H1 (3.0 ± 0.9, *N* = 770, pink). (f) AFM compaction analysis for eNuc-tri alone (99 ± 22 nm, *N* = 260, grey) and after incubation with either eH1.0 (87 ± 16 nm, *N* = 245, blue) or MM-H1 (95 ± 22 nm, *N* = 249, pink). These values represent mean ± s.d. and statistical significance was determined by unpaired two-tailed Student’s *t*-test, **** represents P <0.0001, ns represents P >0.05.

## DISCUSSION

Once believed to be unique to the eukaryotic domain of life, the universe of histone-encoding organisms continues to expand to now include most archaea, some bacteria, and an increasing number of giant viruses (NCLDV). Structural analysis of non-eukaryotic histone-DNA complexes has demonstrated a remarkable diversity in histone-based DNA organization. For example, bacterial histone dimers bind DNA edge-on and filament around straight DNA, while archaeal histone homodimers wrap variable lengths of DNA into dynamic ‘hypernucleosome slinkies’ ^39–41^. In contrast, giant viruses encode clearly recognizable homologs of the four eukaryotic core histones (H2A, H2B, H3, and H4) that assemble into nucleosome-like particles that wrap ∼130 bp of DNA^17,19^. Phylogenetic analysis suggests that these histones diverged prior to the emergence of LECA^20,42–45^. Instances of histone doublets (or even triplets and quadruplets) have been described in NCLDV, which enforce specific pairing of histones and suggest a potential role in the origin of eukaryotic histone dimers/tetramers^2,9,45^. With some NCLDV only encoding one doublet pair, it is possible that NCLDV histones doublets have evolved independent functions in viral genome packaging; although it is unclear if these independent functions were developed before or after complete sets of core histone singlets were encoded in eukaryotes or even MM^44^. MM histones are placed at the root of most histone phylogenetic trees, and thus add to new potential theories regarding nucleosome evolution^5,12,16,20^. Their resemblance to eukaryotic core histones is plausibly a result of acquiring genes through horizontal gene transfer (HGT) post-LECA, however their sequence similarity is low compared to its host *A. castellanii* and even other viral histones. As such, the role of MM core histones in the origin of the nucleosome remains a mystery.

We show that key eukaryotic nucleosome features are conserved within MM-NLPs, despite the low conservation in amino acid sequence. This includes the overall geometry and arrangement of the DNA superhelix, which is achieved through the interactions between the DNA minor groove backbone and the main chain of antiparallel L1-L2 loops of the histones, as well as the N-termini of histone fold α1 helices^1,17,19^. However, MM histones diverge in the lengths of tails and loops within the histone fold which distinctly shape the nucleosomal surface. For example, the MM-H2B loop 1 may act as an ‘arm’ for other interacting viral or host proteins, including putative nucleosome assembly, remodeling factors and histone chaperones, and/or may change the way in which nucleosomes pack against each other into higher order structures. A similar role for an extended loop has been observed for the centromeric histone variant CenpA^46^.

The unique extended MM-H3 C-terminal tail lays across the H3-H4 histone fold. The path of the MM-H3 C-term tail, confirmed by very well-defined density in both the native and crosslinked complexes, encourages interactions with the elongated loops in MM-H2B and promotes unique stabilization of the MM-NLP. While the H3 C-terminal tail is unique to MM, its pathway across H3 α2 mimics that of the Melbournevirus H4 N-term tail, which extends out and lays across the histone fold dimer in the exact same location but from the opposite direction. This conserved requirement of a ‘stabilizing tail’ (either from H3 in MM, or from H4 in Melbournevirus) suggests an ancient feature of the nucleosome that is obsolete in eukaryotic nucleosomes, where histone tails are primarily used for signaling and regulation.

Unlike most histone encoding NCLDV, Medusavirus encodes an acidic putative linker histone H1 that maintains a significant sequence identity to H1 encoded by the host *A. castellanii*. The much higher degree of identity than what is observed for the core MM histones (∼30%) suggests that the predicted H1 was shared through horizontal gene transfer at a later stage. The MM-H1 has the canonical winged-helix domain seen in *X. laevis* H1.0 but only one of the four established residues required for chromatin compaction. The MM-H1 also has an additional acidic winged-helix domain, a feature that is shared with the three linker histones of the host *A. castellanii*. The putative viral linker histone does not compact tri-nucleosomes, suggesting an alternative and independent viral function of the MM-H1.

During viral assembly of Melbournevirus, viral histones reside exclusively in the viral factories of the host cytoplasm to mature into virions upon Melbournevirus infection^17^. In contrast, MM maturation relies heavily on the host nucleus. Viral particles are independently produced in the cytoplasm, but viral DNA replication occurs in the nucleus, and only the assembled capsids that are close to the nuclear membrane are filled with DNA. This could allow the opportunity for the exchange of protein and DNA through open membrane gaps of the MM viral particle positioned near the nuclear membrane, to which MM histones may be used for chromatin organization between or within the host nucleus and the viral particle. Whether the DNA is already assembled into chromatin at the time of packaging, or whether nucleosome assembly takes place in the capsid remains to be determined^47^. The pronounced differences in the four-helix bundle interfaces holding together the (H3-H4)_2_ tetramer, and tethering it to the H2A-H2B dimer, might have evolved to prevent the formation of hybrid nucleosomes consisting of host and virally encoded histones. While histones are essential for Melbournevirus fitness and infectivity, the importance of virally-encoded histones for Medusavirus fitness has yet to be explored^17,20^. As more metagenomes are discovered, more histone-encoding viruses will reveal help fill in the gaps of the evasive evolution of the nucleosome.

## MATERIALS AND METHODS

### Histone sequence alignment and secondary structure prediction

Predicted *Mamonoviridae* family and *Marseilleviridae* family core histone-like protein sequences were aligned with eukaryotic histone sequences with HHpred’s Multiple Alignment using Fast Fourier Transform (MAFFT) with a 1.52 gap open penalty^20,21^. The sequence similarly and identity of *Medusavirus medusae* (MM), *Medusavirus stheno*, Clandestinovirus, Marseillevirus and Melbournevirus relative to each core eukaryotic histone were obtained using the Sequence Manipulation Suite webserver (SMS)^48^. To demonstrate structural conservation of the canonical core histone fold domain between *Eukarya* and *Mamonoviridae*, protein secondary structures were predicted using the MAFFT alignment on the HHpred’s Quick 2D structural prediction webserver^49^. The isoelectric point of each protein was provided by protparam.

Predicted *Mamonoviridae* family and Clandestinovirus linker histone-like protein sequences were aligned against host *Acanthamoeba castellanii* H1*, Xenopus laevis* H1 and *Gallus gallus* H5 with HHpred’s Multiple Alignment using Fast Fourier Transform (MAFFT) with a 1.52 gap open penalty^20,21^. The sequence similarly and identity of MM, Medusavirus stheno and Clandestinovirus, relative to each linker eukaryotic histone were obtained using the Sequence Manipulation Suite webserver (SMS)^48^. To demonstrate structural conservation of the canonical winged-helix domain between *Eukarya* and *Mamonoviridae*, protein secondary structures were predicted using the MAFFT alignment on the HHpred’s Quick 2D webserver^49^.

### MM histone expression, purification, and refolding

MM ORF 318 (H2A), ORF 61 (H2B), ORF 255 (H3), ORF 254 (H4) and ORF 106 (H1) were each cloned into a pET-28a plasmid for expression and purification from *Escherichia coli*. (*E. coli*), with adaptations of well-established eukaryotic histone protocols^24,50^. Expression of each histone was performed in 6 L of Rosetta 2(DE3) *E.coli* cells with induction of 0.5 mM IPTG at OD:0.4-0.6 and growth at 14°C for 18 hrs. Inclusion bodies were isolated from the cells with a minor adaptation from published protocols, including a 30 min stir of the cell suspension in wash buffer containing 50 µL of DNase I, 5 mM MgCL_2_, 50 µg of Lysozyme and 10 mM CaCl_2_ before tissumizing^24^. Isolated inclusion bodies were treated with 1 mL DMSO for 30 min before stirring with 40 mL of the denaturing lysis buffer (6 M Guanidinium HCl, 20 mM sodium acetate pH 5.2 and 200 mM NaCl) for 30 min. After stirring, samples were tissumized for 25 s intervals at 30% approximately 4-5 times until the viscosity of the lysate resembled water. The lysate was spun at 16,000 rpm for 30 min and the supernatant was filtered with a 0.4 µM syringe filter.

The filtered lysate was applied to a 5 mL His-Trap HP column in nickel loading buffer (8 M urea, 20 mM sodium acetate pH 5.2, 200 mM NaCl and 20 mM Imidazole) and eluted utilizing a gradient of the nickel elution buffer (8 M urea, 20 mM sodium acetate pH 5.2, 200 mM NaCl and 1 M Imidazole). Isolated fractions containing each MM core histone (H2A, H2B, H3 and H4) were combined and run over a TSK-SP cation exchange column using the SAUDE 200 buffer (8M urea, 20 mM sodium acetate pH 5.2, 200 mM NaCl, 5 mM βME and 1 mM EDTA) and eluted using a gradient of the SAUDE 1000 buffer (8 M urea, 20 mM sodium acetate pH 5.2, 1000 mM NaCl, 5 mM βME and 1 mM EDTA). For MM H1, fractions from the His-Trap HP column were placed over a Mono-Q anion exchange column in SAUDE buffer. Eluted fractions, from the TSK-SP or Mono-Q, containing each individual histone were combined and dialyzed into 5mM βME for lyophilization. All histones were lyophilized and stored at −20°C^50^. Confirmation of each histone protein ID was performed through LC-MS/MS (Jeremy Balsbaugh, UConn).

For protein refolding, MM H1 was dialyzed alone into refolding buffer (20 mM Tris-HCL pH 7.5, 2 M NaCl, 1 mM EDTA, 1 mM βME) overnight with multiple buffer changes. Precipitated protein was discarded through centrifugation and the supernatant was concentrated for application over a size-exclusion S200 equilibrated in refolding buffer. The sample was stored in 20% glycerol at −80°C. Conversely, MM core histones (H2A, H2B, H3 and H4) were refolded together either as an octamer, as dimers (H2A-H2B), or as putative tetramer (H3-H4)_2_ utilizing previously described protocols^50^. Samples eluted off gel filtration at expected volumes and were stored in 20% glycerol at −80 °C.

### MM-NLP reconstitution

The DNA utilized to reconstitute the MM nucleosome-like particle (NLP) *in vitro* included the Widom 601 DNA at 147 bp (5’– ATCTGAGAATCCGGTGCCGAGGCCGCTCAATTGGTCGTAGACAGCTCTAGCACCGCTTAAACGCACGTACGCGCTGTCCCCCGCGTTTTAACCGCCAAGGGGATTACTCCCTAGTCTCCAGGCACGTGTCAGATATATACATCCGAT –3’), 165 bp (5’– ATCGCCAGGCCTGAGAATCCGGTGCCGAGGCCGCTCAATTGGTCGTAGACAGCTCTAGCACCGCTTAAACGCACGTACGCGCTGTCCCCCGCGTTTTAACCGCCAAGGGGATTACTCCCTAGTCTCCAGGCACGTGTCAGATATATACATCCAGGCCTTGTG GAT –3’) and 207 bp (5’– ATCTAATACTAGGACCCTATACGCGGCCGCATCGGAGAATCCCGGTGCCGAGGCCGCTCAATTGGTCGTAGACAGCTCTAGCACCGCTTAAACGCACGTACGCGCTGTCCCCCGCGTTTTAACCGCCAAGGGGATTACTCCCTAGTCTCCAGGCACGTGTCAGATATATACATCGATTGCATGTGGATCCGAATTCATATTAATGAT –3’) lengths; along with an in-house generated 150 bp 50% G/C content double stranded DNA (5’– GCTAGT CCGTCTTCTACTCTGAAATGAGCAGTCCTAGTCAGCAAGATCGCTCAGCCAACTTTCTACCAGCGCAACCCTAATCTACCCCATGAATGAAGCCGCACCCAAAACCGCATTCTAAGGAGTGACATTAACCCTCGGTGAGGATGT –3’)^25^. MM octamer and DNA were mixed at a range of ratios of DNA to octamer and reconstituted by gradient dialysis (ideal ratio 1.0: 1.6). All eukaryotic histones (*Xenopus laevis*) utilized in reconstitution of eNuc controls were supplied by the Histone Source (Hataichanok Scherman, Colorado State University). Reconstituted samples were analyzed by 5% native-PAGE.

### Sedimentation velocity analytical ultracentrifugation (SV-AUC)

To evaluate the homogeneity, molecular size, and molecular shape of MM-NLPs and MM-LE-tri-NLP in solution; we used SV-AUC with absorbance optics (λ = 280 nM). MM-NLP of various DNA lengths and MM-tri-NLP, all at 250 nM, were spun at 30,000 – 35,000 rpm at 20°C in the Beckman XL-A ultracentrifuge using the An60Ti rotor (50 mM NaCl, 20 mM Tris-HCL, 1 mM EDTA, pH 7.5 and 1 mM DTT). Sedimentation velocity data analysis were performed in UltraScan III version 4.0 to determine sedimentation coefficients (S _(20,W)_), frictional ratios (f/f_0_) and molecular weights of each sample using established protocols ^17,51,52^. Integral S _(20,W)_ distributions plots are displayed using GraphPad Prism version 10.0.0 for Windows, GraphPad Software, Boston, Massachusetts USA, www.graphpad.com.

### Thermal Stability Assay

MM-NLPs and eNucs were reconstituted on Widom 601 DNA (207 bp) and Random 150 bp DNA as described above. Melbournevirus NLPs were also reconstituted on Widom 601 at 207 bp DNA as previously described^17^. Two replicates of each nucleosome (800 nM) were incubated in the presence of SYPRO Orange for 1 min at 25 °C, in 20 mM Tris-HCl (pH 7.5), containing 5 mM DTT and 50 mM NaCl. SYPRO Orange is provided SIGMA-ALDRICH as a 5000x concentrated solution (Catalog #: S5692). Each reaction was performed in a final volume of 20 µL, containing 2.5 µL of 62.5-fold diluted SYPRO Orange 5000x (final concentration 8x).

Samples were increased by 1 °C and maintained for 1 min at the increased temperature. Fluorescence measurements were measured using a StepOnePlus Real-Time PCR unit (Applied Biosystems) after each 1 min incubation step. This sequential process was repeated for each 1 °C from 25 °C to 95 °C using the Cal Orange filter (fluorescence emission maximum, 560 nm).

Normalized plots displaying the relative fluorescence release of SYPRO Orange over 25 °C to 95 °C are displayed using GraphPad Prism version 10.0.0 for Windows, GraphPad Software, Boston, Massachusetts USA, www.graphpad.com. The thermal melting point (Tm) ranges of each NLP sample (n=2) measured in the thermal stability assay were determined by locating the temperature at the lowest point of the fluorescence derivative (-d/dU RFU).

### Sucrose gradient sedimentation and gradient fixation (GraFix) crosslinking

A continuous 10-30% (w/v) sucrose gradient was prepared in a 13.2 mL Beckman polypropylene coulter tube (331372). To form the gradient, 6 mL of the top solution (50 mM NaCl, 20 mM HEPES, 1 mM EDTA, pH 7.5 and 10% sucrose) was added to the coulter tube and then 6 mL of the bottom solution (50 mM NaCl, 20 mM HEPES, 1 mM EDTA, pH 7.5 and 30% sucrose) was slowly added from the bottom of the tube, pushing the top solution up until the demarcation line reached halfway. A gradient maker (BioComp Gradient Master) was then used to generate the continuous gradient. A 200 µL MM-NLP sample (3 µM) was loaded on top of the gradient and spun at 4°C for 18 hr at 30,000 rpm (Beckman, Rotor SW-41Ti). Samples were fractionated with absorbance optics (λ = 260 nm) for identification of complexes (BioComp Gradient Master). Desired complex fractions were determined using 5% native-PAGE and SDS-PAGE to be combined and dialyzed to remove sucrose (50 mM NaCl, 20 mM Tri-HCL, 1 mM EDTA, pH 7.5 and 1 mM DTT). Gradient fixation (GraFix) was performed utilizing the same procedure, with the addition 0.15% glutaraldehyde in the bottom solution to form a continuous gradient with crosslinker. After centrifugation, fractions were dialyzed to remove sucrose and quench the crosslinking reaction (50 mM NaCl, 20 mM Tri-HCL, 1 mM EDTA, pH 7.5 and 1 mM DTT).

### Homology modeling

Initial homology modeling of the MM histones ORF 318 (H2A), ORF 61 (H2B), ORF 255 (H3) and ORF 254 (H4) were constructed using SWISSMODEL, with *Xenopus laevis* histones in the context of a nucleosome (1AOI) as a reference^53–55^. To identify steric clashes in the initial model, CPPTRAJ of the Amber MD package (v18) with a cutoff distance of 0.8 Å between over-lapping atoms was utilized^56^. Each clash was manually addressed by modifying rotamers to reduce overlap and to achieve energy minimization in Chimera using default settings^57–59^. This initial MM nucleosome conformation was further refined by fitting into final 3D electron maps as described below.

### Single particle cryo electron microscopy (cryo-EM) and data processing

MM-NLP particles isolated from both sucrose gradient sedimentation (native) and GraFix (crosslinked) were concentrated to 2 µM using Amicon Ultra-4 centrifugal filters (Ultracel 30K, Millipore). C-Flat 1.2/1.3 (Cu) grids were glow discharged (Tergeo-EM Plasma Cleaner) at 40 mA for 30 s before 4 µL of sample was applied to the grid and plunged into ethane using the Vitrobot Mark IV at 100% humidity at 4°C with no wait time, a 2 s blot, and a blot force of 5. Micrographs of the MM native and crosslinked nucleosome particles were acquired with a nominal magnification of 64000x on a GEI Titan Krios (300 kV) outfitted with a Gatan K3 direct detection camera. The raw pixel size was 1.017 Å with movies captured in super-resolution mode maintaining an electron dose rate of 46.29 e/Å^2^. The defocus range was −0.8 to −2.2 µM.

Both datasets were processed initially by cryoSPARC (v2.12.4) through motion correction and CTF estimation^60,61^. Exposures were curated to exclude sub-optimal characteristics by inspecting the CTF Fit resolution (Å), relative ice thickness and defocus range. Approximately 1000 particles were manually picked from the curated exposures to generate a picking model through Topaz (downsampling=16). For the crosslinked MM-NLP, Topaz picking yielded 451,081 particles which were subjected to two iterative rounds of 2D classification in order to discard bad particles^62^. Selected particles (162,284) were used to generate three *ab initio* models and the 3D model with the best directional distribution was selected to improve to the final resolution using homogeneous refinement, non-uniform refinement and local refinement (154,200 particles) (**Figure S3B**)^63,64^. For the native MM-NLP, Topaz picking yielded 72,768 particles and these particles were also subjected to two iterative rounds of 2D classification to remove undesirable particles. This yielded 72,768 particles which were utilized to generate *an initio* model that was similarly subjected to homogeneous refinement, non-uniform refinement, and local refinement to improve resolution (**Figure S3A**).

### Structural characterization of MM nucleosomes

Comparisons between the MM nucleosome (8UA7), Melbournevirus nucleosome (7N8N) and eukaryotic nucleosome (1AOI) were conducted using ChimeraX^65–67^. Structures were aligned with the matchmaker tool in ChimeraX using the best-aligning pair of protein chains between the reference and matching structure. All residue and feature comparisons were determined by previously identified canonical eukaryotic histones features and figures were rendered with ChimeraX^1,65^.

### Structural prediction of putative linker histone MM-H1

The sequence of the putative linker histone MM-H1 was folded using AlphaFold (v.2.3.2)^68^. The highest ranked model was then aligned to the *X. laevis* linker histone H1 in PDB 5NL0 using ChimeraX.

### MM-tri-NLP reconstitution

Tri-NLP reconstitution was performed using the Linker Ended Widom 601-207 bp x3 repeat DNA (LE-Tri) (5’– ATCTAATACTAGGACCCTATACGCGGCCGCATCGGAGAATCCCGGTGCCGAGGCCGCTCAATTGGTCGTAGACAGCTCTAGCACCGCTTAAACGCACGTACGCGCTGTCCCCCGCGTTTTAACCGCCAAGGGGATTACTCCCTAGTCTCCAGGCACGTGTCAGATATATACATCGATTGCATGTGGATCCGAATTCATATTAATCATATCTAATACTAGGACCCTATACGCGGCCGCATCGGAGAATCCCGGTGCCGAGGCCGCTCAATTGGTCGTAGACAGCTCTAGCACCGCTTAAACGCACGTACGCGCTGTCCCCCGCGTTTTAACCGCCAAGGGGATTACTCCCTAGTCTCCAGGCACGTGTCAGATATATACATCGATTGCATGTGGATCCGAATTCATATTAATCATATCTAATACTAGGACCCTATACGCGGCCGCATCGGAGAATCCCGGTGCCGAGGCCGCTCAATTGGTCGTAGACAGCTCTAGCACCGCTTAAAC GCACGTACGCGCTGTCCCCCGCGTTTTAACCGCCAAGGGGATTACTCCCTAGTCTCCAGGCACGTGTCAGATATATACATCGATTGCATGTGGATCCGAATTCATATTAATGAT –3’), and an in-house generated 500 bp 50% G/C content double stranded DNA (5’– GCTAGTCCGT CTTCTACTCTGAAATGAGCAGTCCTAGTCAGCAAGATCGCTCAGCCAACTTTCTACCAGCGCAACCCTAATCTACCCCATGAATGAAGCCGCACCCAAAACCGCATTCTAAGGAGTGACATTAACCCTCGGTGAGGATGTCCATACAAGCACCTCCTACTACGGATCGAACCGTTAGTTCCCCAACTAAGTCCAAACCGTTAGACCGCTTTCCGTACCATTCCGGTACTTATCTTCGCCACAACCTGAGACAATCCCAAGCTTAAGGCTCGACACAGACTGACGAAGGATATATCTCGCCCTAACCGTACCTCTATACCGCCATGAAGGAAGTGCCAAGTAGCCACAGAACCTTGGGATAGCAAGACTCTATGTCCCAGACCTCACTAACACCGAAGGAAAGTACCCACACAGACATCAGGAAAACCCTCTGACCACTACGGCGAATGAAAAGTCCAGAGGACCAATACGTTACAGAGGCGACTGGATGT –3’)^25^. MM octamer and DNA were mixed at a 1.0: 4.8 ratio of DNA to octamer and reconstituted into tri-nucleosomes by gradient dialysis^24^. All eukaryotic histones (*Xenopus laevis*) utilized in reconstitution of LE-tri controls were supplied by the Histone Source (Hataichanok Scherman, Colorado State University). Reconstituted samples were tested for quality using 4% native-PAGE and mass photometry to confirm a homogenous sample.

### Mass Photometry

Mass photometry measurements were performed on a Refeyn TwoMP mass photometer (Refeyn Ltd). Glass coverslips were first cleaned with isopropanol, deionized water, and dried with N_2_ gas, before coating with a 0.01% Poly-L-Lysine solution for 20 seconds, rinsing with water, and drying with N_2_ gas. To form a sample chamber, self-adhesive silicon gaskets were adhered to the top of the treated coverslip. For each measurement, the coverslip was placed on the oil-immersion objective lens, centered on a single well, and 13.5 µl sample buffer (20 mM HEPES, pH 7.5, 100 mM KCl) was added to the well and the focal position of the glass surface was determined and held constant using an autofocus system. Nucleosome samples were first diluted to 100-200 nM, before a final 10-fold dilution onto the sample stage (final concentration of 10-20 nM). All dilutions were performed at room temperature in sample buffer (20 mM HEPES, pH 7.5, 100 mM KCl). A 60 second video was recorded immediately after the final dilution. A fresh well and dilution was used for each measurement and repeated at least three times for each sample. Tri-nucleosomes were diluted in buffer so that the number of detected events (particle counts) during the 60 second measurement was roughly 4,000 to 9,000 for optimum data acquisition and processing. A known mass standard (β-amylase and thyroglobulin) was used to convert image contrast-signal into mass units. To calculate the molecular weight of the main species observed on the particle counts versus molecular mass distribution histograms we used the Gaussian function in the DiscoverMP software.

### Atomic Force Microscopy (AFM)

MM-tri-NLP and eNuc-tri were reconstituted via salt gradient dialysis as previously described^37^. AFM slides were prepared by freshly cleaving the mica, then treated with APTES for 30 minutes, rinsed with water, and dried with nitrogen gas utilizing a 0.22 µm PES filter. Tri-nucleosomes samples were diluted in 20 mM Tris-HCl pH 7.5 and 1 mM EDTA to a final concentration of 1 or 2 ng/µl. Immediately after dilution, samples were applied to the APTES mica slide for 2 minutes, rinsed with water, and dried with filtered N_2_ gas. For H1 samples, the respective tri-nucleosome was incubated with H1 in a 1:4 ratio for 30 minutes at room temperature, before being diluted 2-fold and deposited on the APTES surface (as described above). All samples were imaged in air on JPK/Bruker NanoWizard 3.0 with TAP300-Gold (Ted Pella) cantilevers. Images were collected at a scan size of 1×1 or 2×2 µm, a resolution of 512 × 512 pixels, and at 1-3 Hz.

AFM data was leveled, processed, and analyzed using the Gwyddion software^69^. For particle height analysis, a mask at 1.5 nm was applied to identify particles, followed by extraction of the maximum height for each particle. Rare particles (>8 nm) were excluded as debris or aggregation. For compaction analysis, total distance between nucleosomes was measured in Image J by manually measuring the perimeter of a triangle formed between the three nucleosomes. Of note, only tri-nucleosomes where three nucleosomes were visible were included in this analysis. Analysis and graphing of extracted data was completed in GraphPad Prism (GraphPad Software, Boston, Massachusetts USA, www.graphpad.com).

### Circular Dichroism (CD)

Far-UV CD spectra of H1 proteins (0.5 mg/ml) were recorded at room temperature on a ChirascanPlus (Applied Photophysics Ltd, UK) spectrometer using quartz tubes (0.5 mm optical path length). The measurements were recorded in the 180-260 nm wavelength range with a 0.5 nm step size. All experiments were carried out in 0.02 M Sodium Phosphate and 0.2 M Sodium Fluoride at pH 7.5. For each H1 protein, five replicate CD spectra were averaged, baseline-corrected for signal contributions by the buffer. For secondary structure analysis of CD spectra, the DichroIDP program was used^70^. This CD analysis program was chosen as it is suitable for analyses of proteins containing significant amounts of disordered structures (as seen in eH1.0). Data was graphed using Graph Pad Prism (GraphPad Software, Boston, Massachusetts USA, www.graphpad.com).

## Supporting information

all supplementary info

## ACKNOWLEDGEMENTS

We thank Charles Moe (previously CU Boulder) for help with data collection at the BioKEM facility and Dr. Annette Erbse for training and assistance in the Shared Instrument Facility at CU Boulder. We thank Dr. Shawn Laursen for the design of the 50% G/C 150 bp sequence DNA utilized in thermal assays. NMH is a Howard Hughes Medical Institute Fellow of the Damon Runyon Cancer Research Foundation, DRG-2499-23.

## AUTHOR CONTRIBUTIONS

CMT – Conceptualization; Methodology; Investigation; Data curation; Validation; Formal Analysis; Writing (Original Draft, Review and Editing); Visualization

NMH – Investigation; Data curation; Validation; Formal Analysis; Writing (Review and Editing); Visualization

KL – Conceptualization; Funding Acquisition; Project Administration; Supervision; Writing (Original Draft, Review and Editing)

## DATA AVAILABILITY

Structure data were deposited at the appropriate repositories (PDB entry ID 8UA7; EMDB entry ID EMD-42053). All other data will be made available upon publication.

