## Supplementary material for "Characterization of *Medusavirus* encoded histones reveals nucleosome-like structures and a unique linker histone": all supplementary info

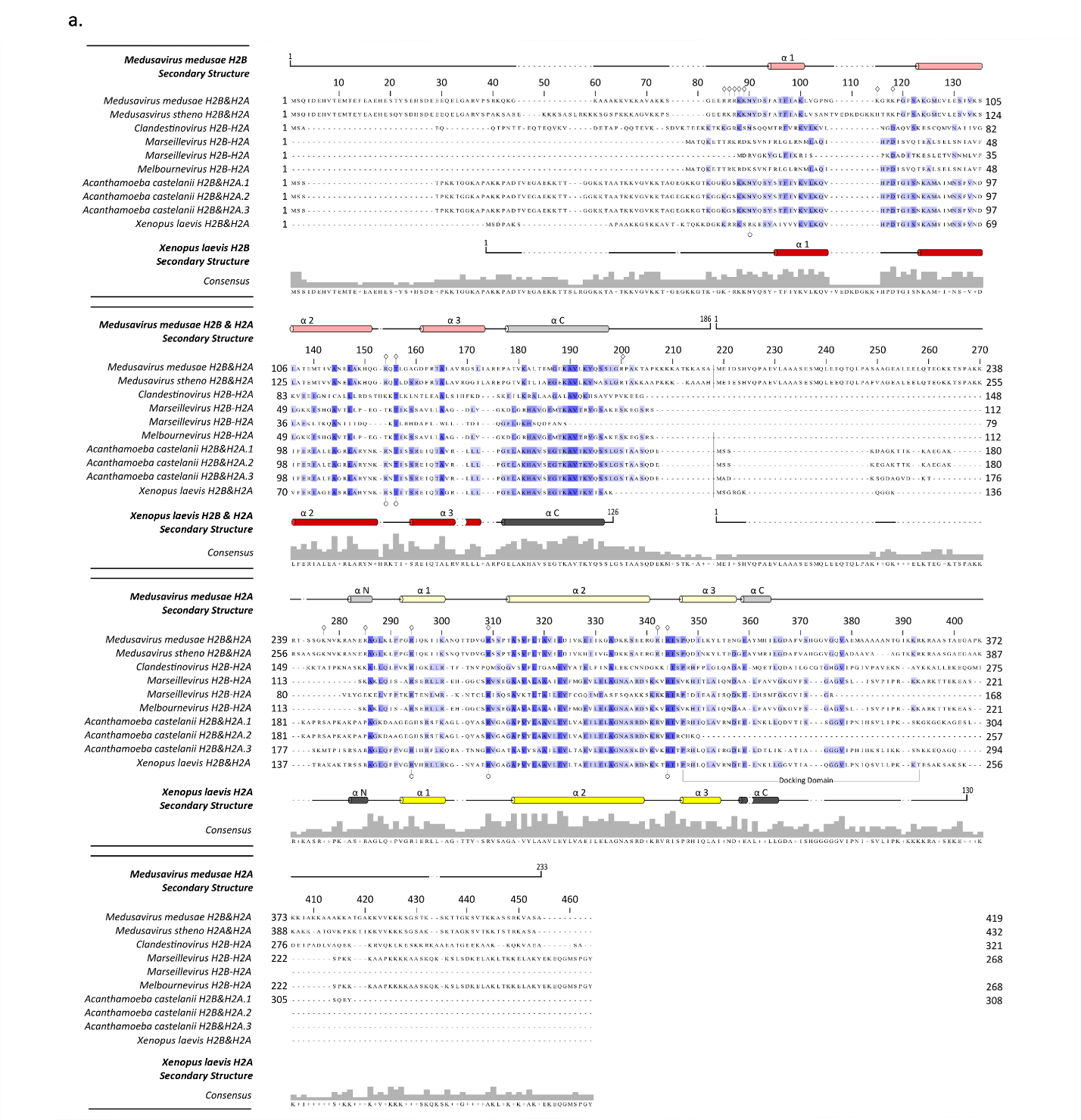

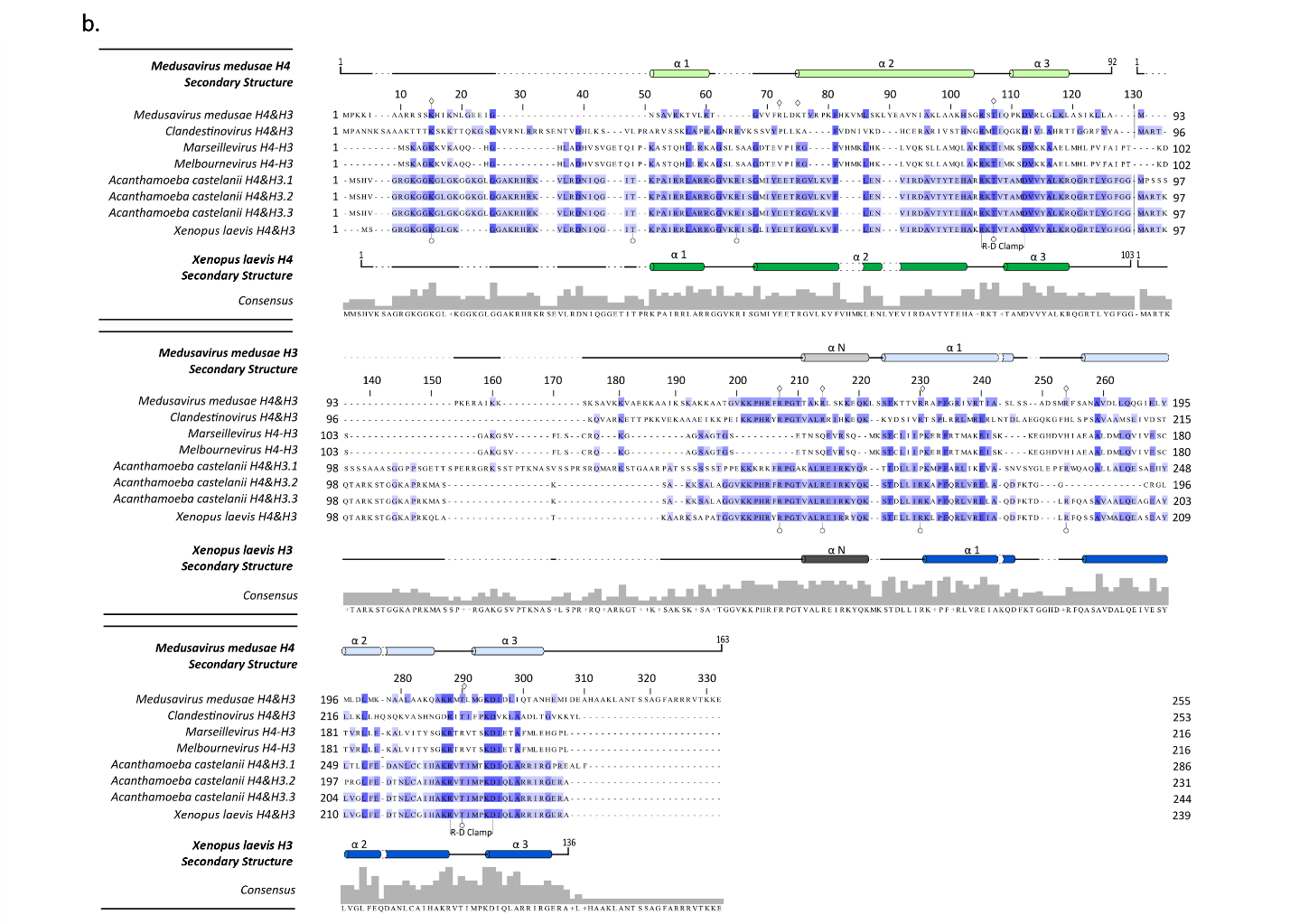

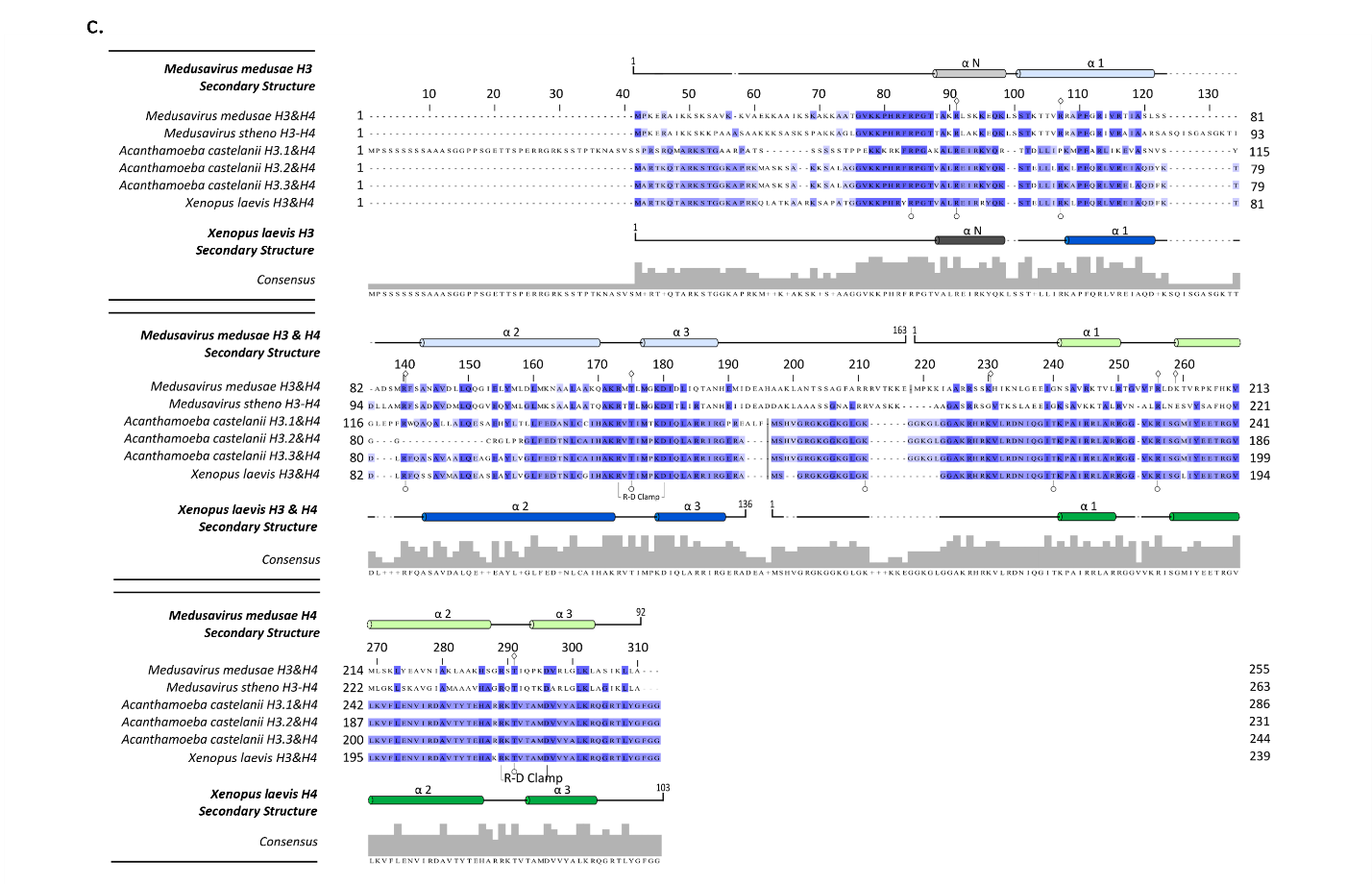

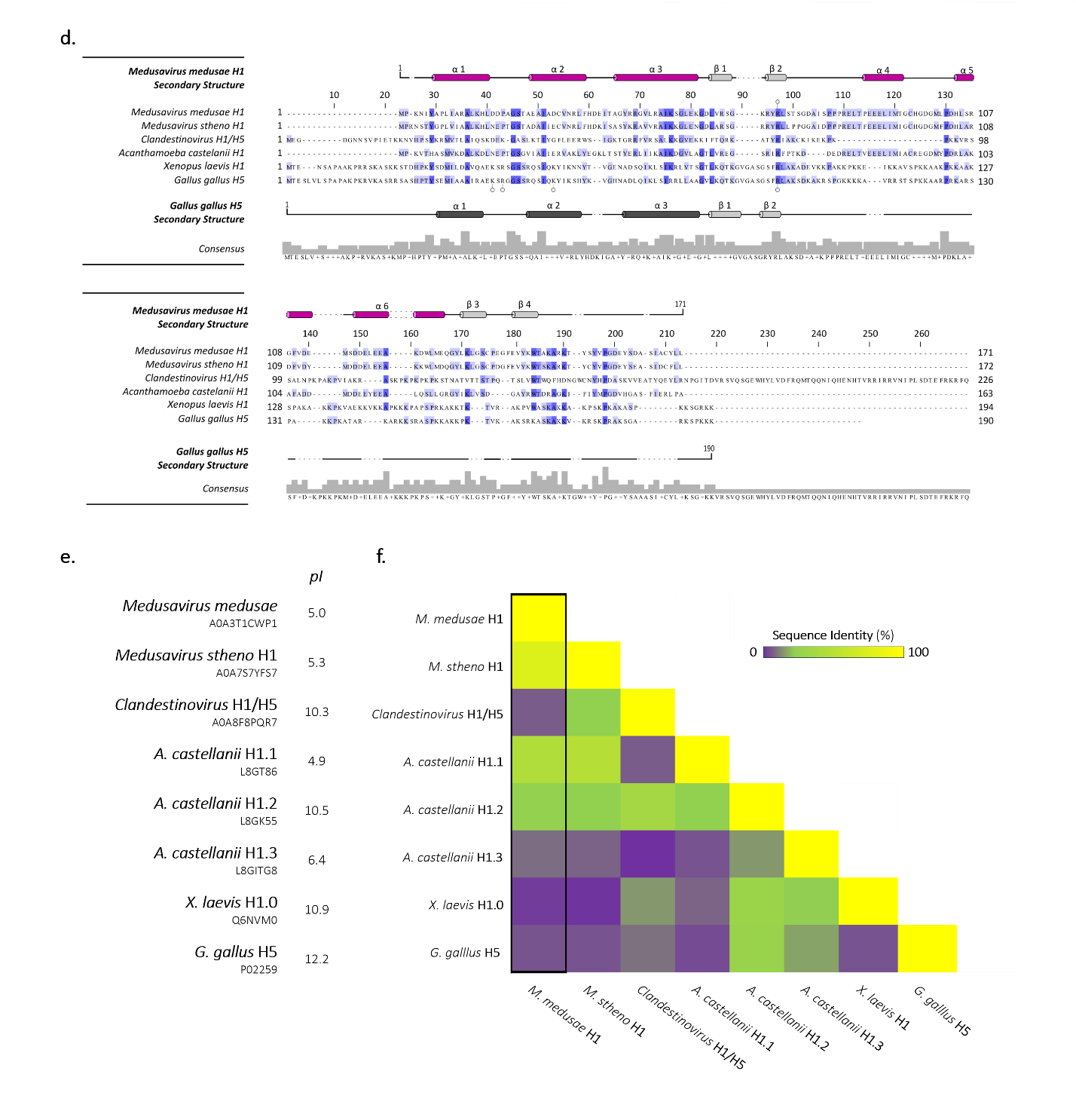

**Figure S1. Complete sequence alignment and secondary structure prediction of *Medusavirus* *medusae* histones.**

Viral histone dimer pairs (or doublets) H2B-H2A and H4-H3 were aligned against *A. castellanii* and *X. laevis* histones using HHPRED’s multiple sequence alignment tool, ClustalΩ. Conservation of each residue within the alignment is represented by blue shading, where darker blue signifies a greater conservation. Known α helices of *X. laevis* are shown in dark colored tubes (H2B-red, H2A-yellow, H4-green, H3-blue). Predicted α helices of MM (light colored tubes) were generated using HHPRED’s Quick 2D prediction web server. (a) Complete sequence alignment of H2B and H2A viral histones (*Mamonoviridae* and *Marseilleviridae* families) against viral host histones *A. castellanii,* and *X.laevis*. (b) Complete sequence alignment of H4 and H3 viral histones (excluding *M. stheno* doublet) against viral host *A. castellanii*, and *X, laevis*. (c) Complete sequence alignment of H3 and H4 viral histones (including *Medusavirus stheno* H3-H4 doublet) against viral host *A. castellanii*, and *X. laevis*. This differs from previous alignment in order of histone pairs (H3-H4 instead of H4-H3). (d) Viral putative linker histone H1 was aligned against *A. castellanii, X. laevis*, and *Gallus gallus* H5. Known α helices of *X. laevis* H1 are shown in dark grey colored tubes. (e) Isoelectric point (pI) of each predicted and known linker histone H1. (f) Heat map comparing percent identity of predicted viral linker histone H1/H5 and eukaryotic H1/H5 sequences. MM-putative H1 is outlined in black. Related to Figure 1.

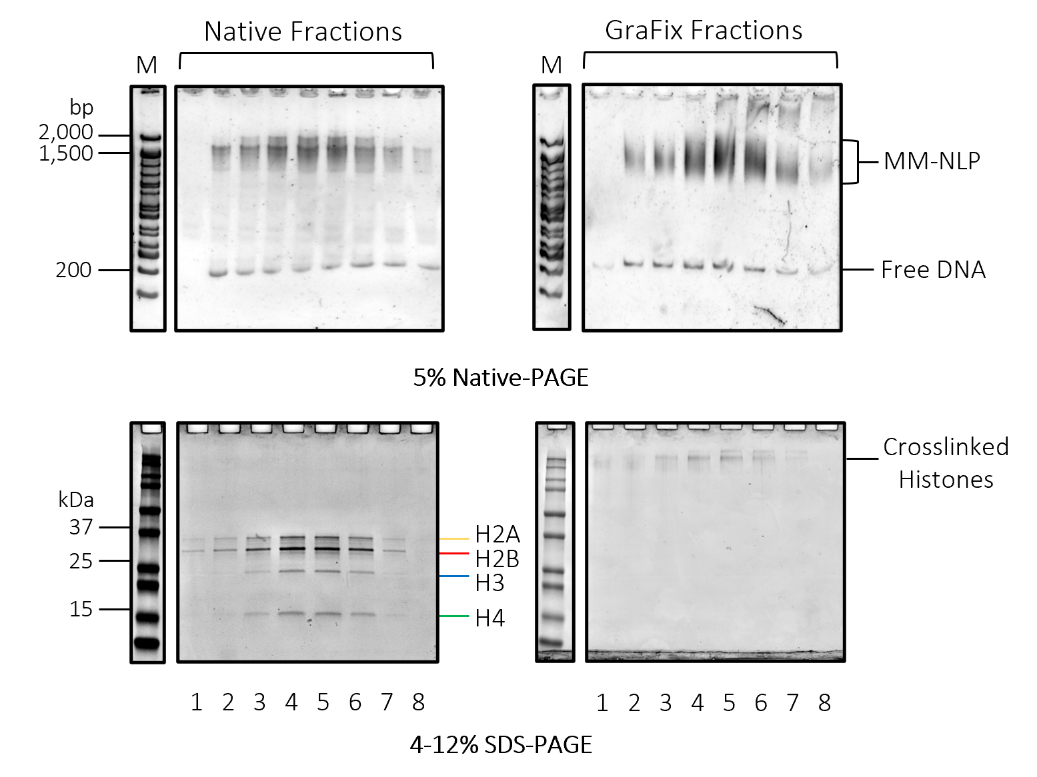

**Figure S2. MM-NLP preparation for Cryo-EM.**

Sucrose gradient sedimentation and GraFix of MM-NLP with 207 bp DNA (MM-NLP_207W_). Fractions of each were analyzed by 4-12% SDS-PAGE stained with BlazinBlue (protein visualization) and 5% Native-PAGE stained with SYBRGold (DNA visualization) to determine composition of particles. Related to Figure 2 and 3.

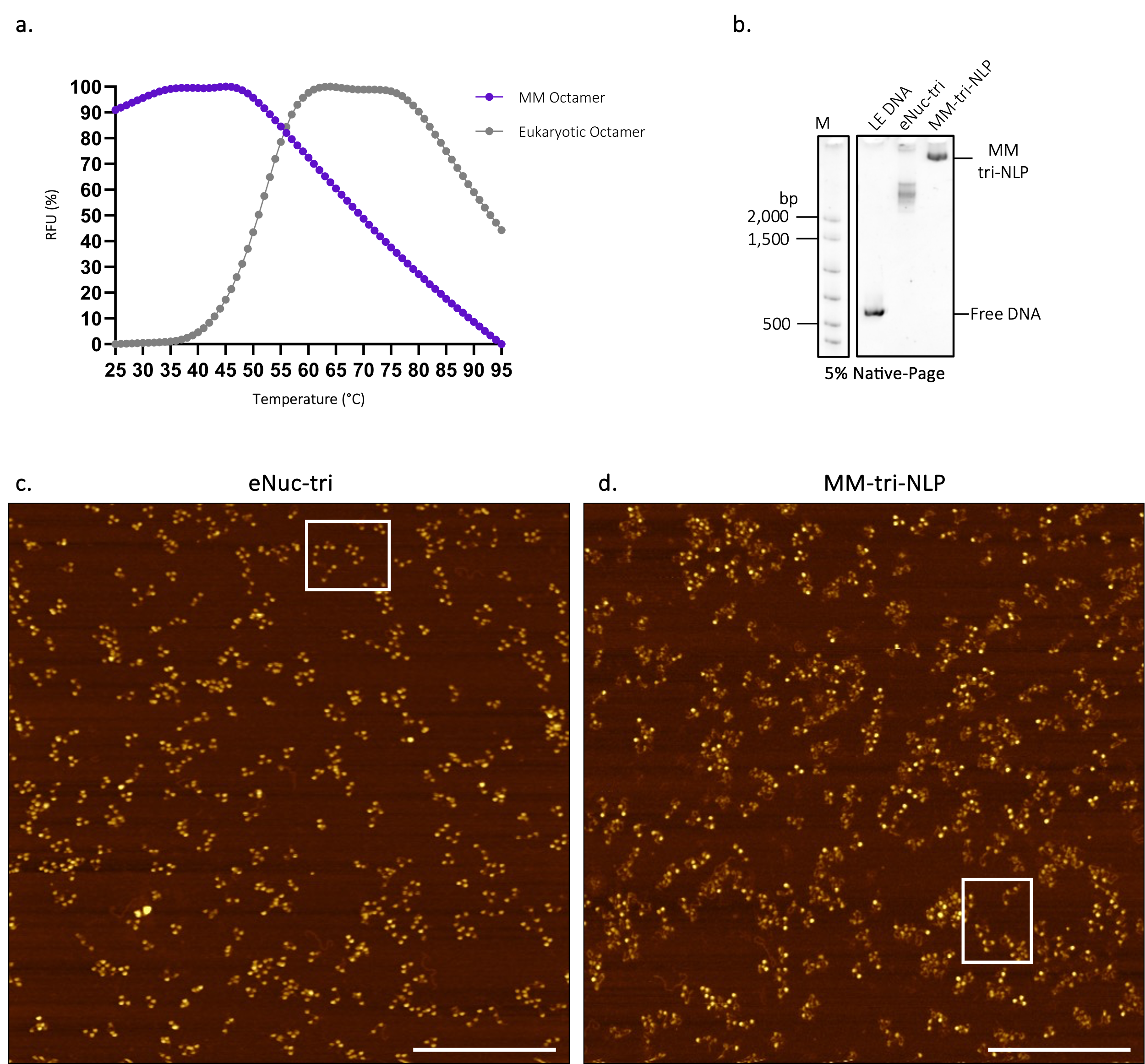

**Figure S3. Biochemical analysis of *Medusa medusae* octamers and tri-nucleosomes.**

(a) Thermal shift stability of MM and eukaryotic octamer utilized in formation of NLP. The raw relative fluorescence units were normalized for plotting. (b) MM tri-nucleosomes (MM-tri-NLP) and eukaryotic tri-nucleosomes (eNuc-tri) on LE DNA. Representative AFM topography images of (c) eNuc-tri and (d) MM-tri-NLP, white squares represent particles shown in Figure 2g. Scale bar = 500 nm. Related to Figure 2.

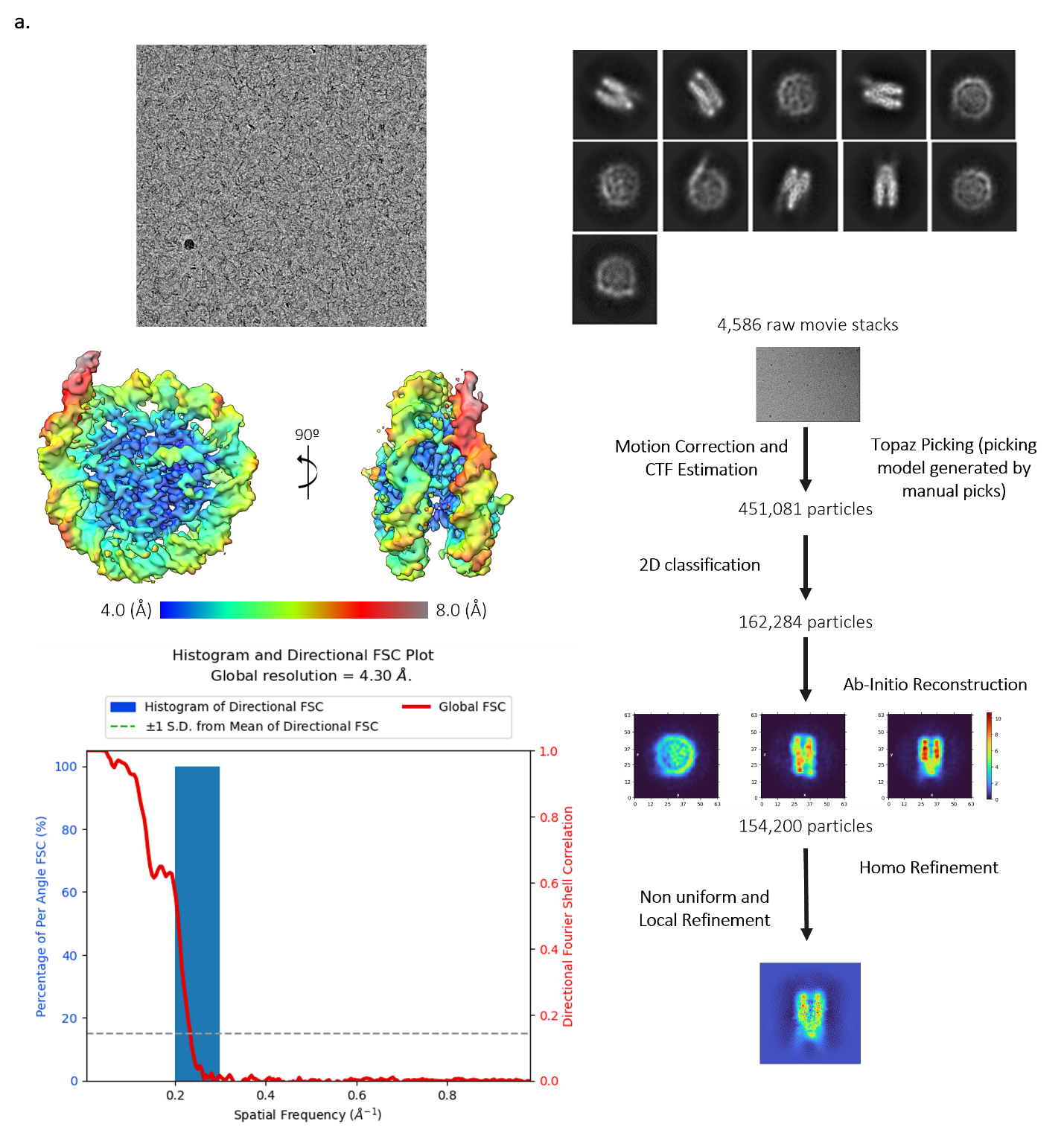

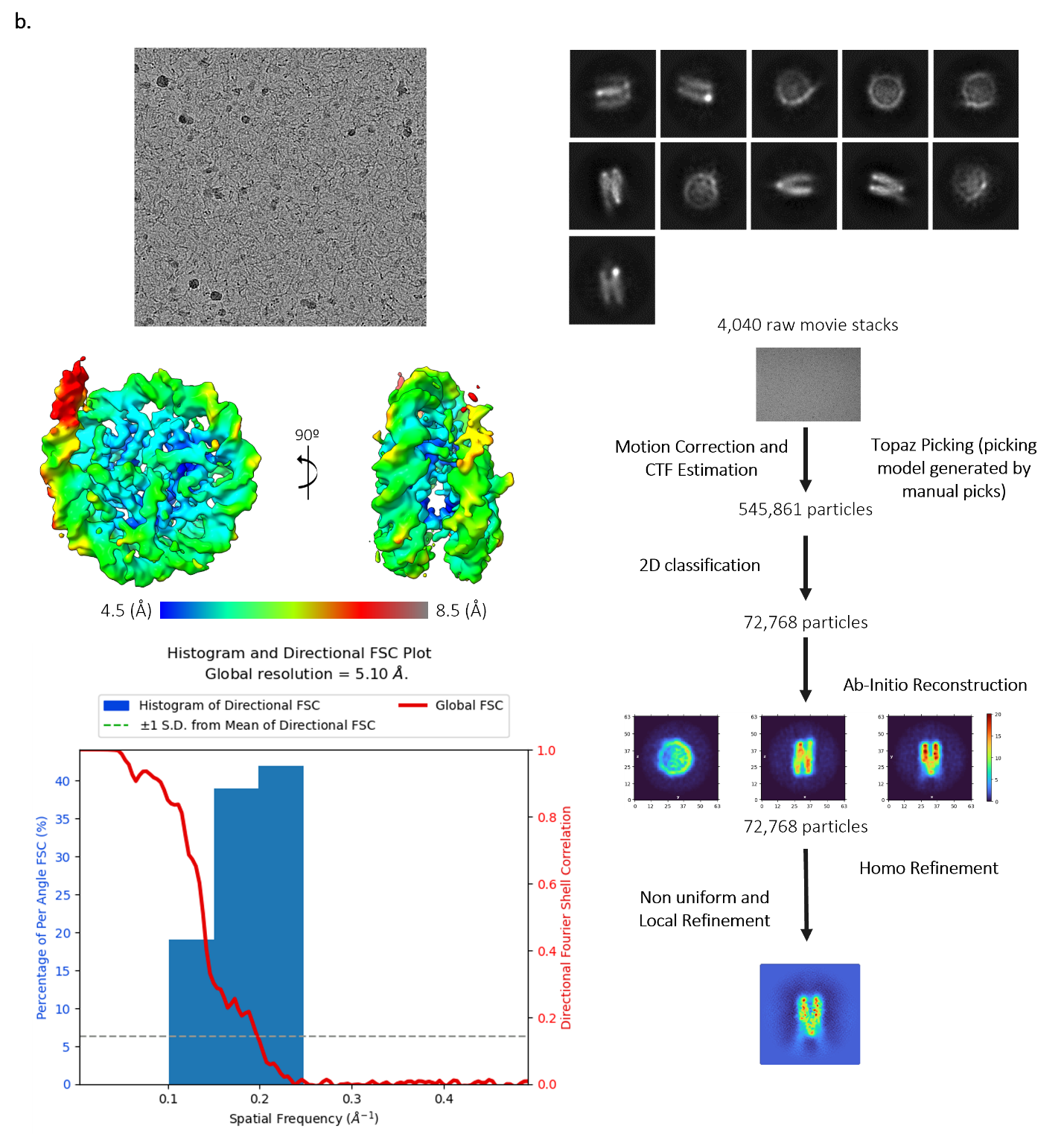

**Figure S4. Cryo-EM analysis of Native and GraFix MM-NLP_207 bp_ .**

(a) Raw micrograph of GraFix MM-NLP_207_, 2D class averages generated from represented dataset, 3D structure of MM-NLP with local resolution map, FSC curve, and CryoSPARC data processing flow chart. (b) Raw micrograph of native MM-NLP_207_, 2D class averages generated from represented dataset, 3D structure of MM-NLP with local resolution map, FSC curve, and CryoSPARC data processing flow chart. Related to Figures 3, 4 and 5.

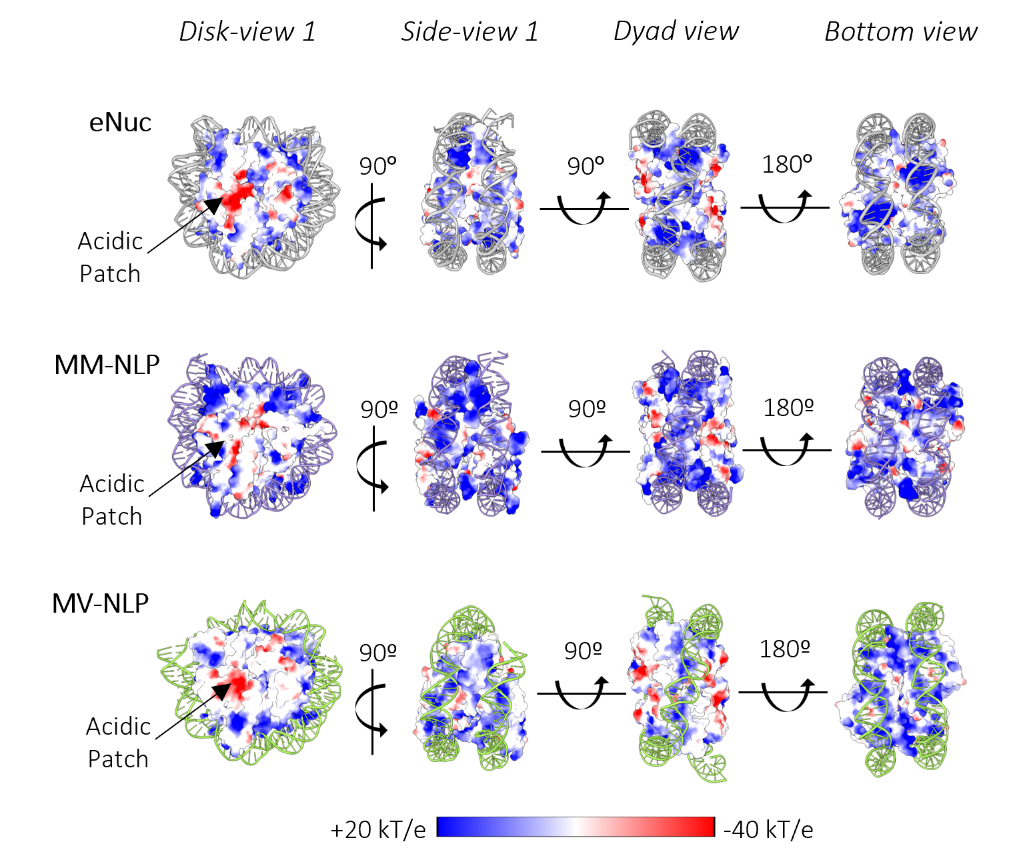

**Figure S5. Electrostatic surface representation comparison of eNuc to viral NLPs.**

Charged surface representation of histones from the eNuc (PDB ID: 3LZ0), MM-NLP and Melbournevirus-NLP (MV-NLP; PDB ID: 7N8N) in different orientations. Related to Figure 5.

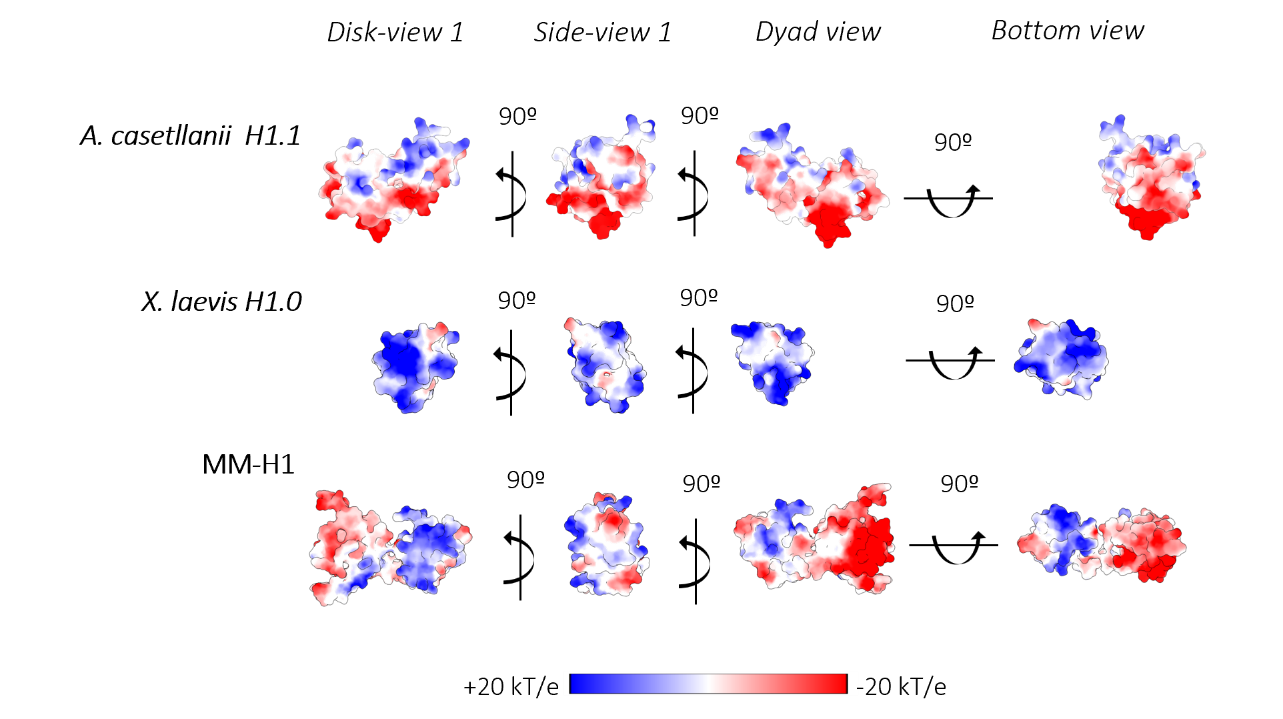

**Figure S6. Electrostatic surface representation comparison of host *A. castellanii* H1.1 and *X. laevis* H1.0 to *Medusa medusae* linker histone H1.**

Charged surface representation of *Acanthamoeba castellanii* H1.1, *Xenopus laevis* H1.0, and MM-H1 with rotational views. Coordinates for *Xenopus laevis* H1 were acquired from 5NL0. Coordinates for *Acanthamoeba castellanii* H1.1 and MM-H1 were determined through AlphaFold. Related to Figure 6.

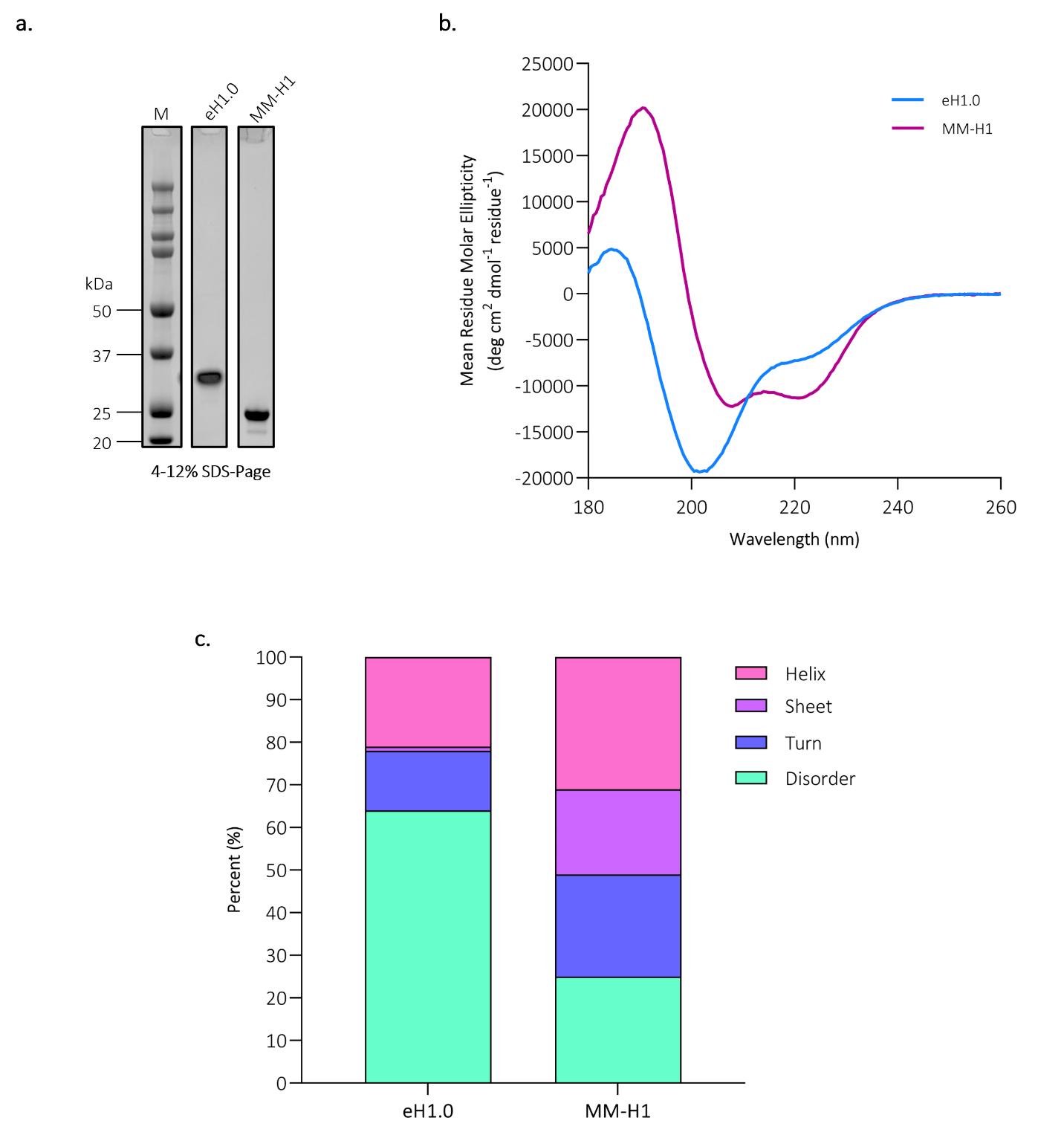

**Figure S7. Biochemical analysis of *Mus musculus* and *Medusavirus medusae* histone H1.**

(a) Purified MM-putative linker histone H1 (MM-H1) and *Mus musculus* H1.0 (eH1.0). (b) CD spectra of purified eH1.0 (blue) and MM-H1 (pink). (c) Secondary structure estimation based on experimental CD data (shown in b), using DichroIDP. Related to Figure 7.
